# Aging promotes lung cancer metastasis through epigenetic ATF4 induction

**DOI:** 10.1101/2024.07.03.601209

**Authors:** Angana A.H. Patel, Jozefina Dzanan, Kevin X. Ali, Ella A. Eklund, Samantha W. Alvarez, Ilayda Altinönder, Dorota Raj, Ahmed Ezat El Zowalaty, Sama I. Sayin, Martin Dankis, Maria Schwarz, Emma Jonasson, Kristell Le Gal, Heba Albatrok, Roger Olofsson Bagge, Anetta Härtlova, Anders Ståhlberg, Andreas Hallqvist, Clotilde Wiel, Volkan I. Sayin

## Abstract

Lung cancer is primarily a disease of the elderly. Despite shared molecular changes between aging and cancer ^1^ – such as permissive chromatin states and deregulated protein homeostasis – studies on physiologically aged models of human lung cancer are lacking. Here, we show that aging alters the progression of KRAS-driven non-small cell lung cancer (NSCLC), promoting metastasis while suppressing primary lung tumor growth. Clinically, a multicenter analysis of all consecutively diagnosed NSCLC cases in Western Sweden over a 3-year period confirmed increased metastasis and smaller primary tumor size with age in KRAS-driven NSCLC. In addition, primary lung tumor cultures derived from older mice demonstrated an increased metastatic phenotype. Unbiased transcriptomic and epigenomic analyses identified ATF4, a major arm of the unfolded protein response (UPR), as a driver of aging-induced lung cancer metastasis. Furthermore, we found that the age-associated increase in ATF4 fuels metastatic dissemination through metabolic rewiring, including increased glutaminolysis. Finally, we report that pharmacological inhibition of glutaminase effectively suppressed aging-induced metastasis. Our findings suggest a novel adjuvant therapy for human lung cancer by targeting aging-induced metabolic plasticity, highlighting the need to consider the biology of aging in the development of cancer therapy.

## Introduction

Aging and cancer are complex processes that share many molecular and cellular changes, including disrupted protein homeostasis, epigenetic remodelling, and altered nutrient sensing, suggesting an intricate mechanistic relationship between the two^1^. However, no study has directly investigated the contribution of aging-induced epigenetic alterations on lung cancer progression. Little is known about the impact of aging on lung cancer development^2^, especially in genetically engineered mouse models (GEMM) of cancer, which are pivotal for dissecting the biology of lung cancer and drug discovery^3^.

Lung cancer remains the leading cause of cancer-related deaths worldwide, with a 5-year survival rate of only around 20%^4^. Metastatic disease remains the primary driver of mortality among lung cancer patients, despite significant progress made in early detection, targeted-, and immunotherapies. Diagnosed at a median age of 70, with a peak incidence in the age group between 65-75 years, lung cancer predominantly affects the elderly, with age significantly influencing both incidence and outcomes^5^.

Age-associated changes in humans are largely conserved in mice, making them an excellent model for studying mammalian aging^6^. Aging-associated cellular and molecular alterations conserved between mice and humans start to emerge in mice at 18-19 months of age, which 5-6 months later – the typical duration of a lung cancer experiment– corresponds to human age of 70 years^7,8^. GEMMs such as the KP model of non-small cell lung cancer (NSCLC) are indispensable for studying the effects of aging on cancer progression, as they develop tumors in a natural microenvironment and recapitulate critical features of human tumors^3^. Despite their potential, these models are predominantly utilized in studies involving young mice, leaving a gap in our understanding of age-related NSCLC pathogenesis.

Emerging evidence indicates that age-related alterations in the tumor microenvironment can drive metastatic progression, as demonstrated in melanoma models^9–11^. However, the specific and cell-intrinsic impact of age-related changes on lung cancer progression and metastasis remains unknown. Importantly, no study has yet explored the impact of physiological aging on metastasis using GEMMs of lung cancer. Here we use young and old KP-mice to define the impact of physiological aging on lung cancer progression and metastasis.

## Results

### Aging alters lung cancer progression

To investigate the impact of aging on lung tumorigenesis, we simultaneously induced lung tumors in ***K****ras*^LSL-G12D/-^; *Tr**p***53^flox/flox^ (*KP*) mice aged 2-3 months (*KP*-Young) and 18-19 months (*KP*-Old) via intratracheal instillation of Cre-encoding viral particles (Fig. 1a). These age groups were chosen to represent two distinct stages: early adulthood, which is the typical age used in lung cancer studies^3^, and an older age where molecular aging phenotypes become relevant^4,7,8^. The old group corresponds to the median age at diagnosis for NSCLC at 70 years^4^, accurately reflecting the demographic most affected by NSCLC (Extended Data Fig. 1a). Surprisingly, we observed decreased primary lung tumor burden in *KP*-Old mice compared to *KP*-Young mice (Fig. 1b). Immunohistochemical analysis revealed decreased proliferation in *KP*-Old tumors, as judged by phosphorylated-histone 3 positive cells, while all tumors from *KP*-Old and *KP*-Young mice were positive for the alveolar type 2 (AT2) lineage marker pro-SPC (Extended Data Fig. 1b and 1c), the main cell type of origin for lung adenocarcinoma. To rule out unequal tumor initiation in the aging lung, we next induced lung tumors in young and old *KP* mice with a viral vector driven by a SPC promotor (Adeno-SPC-Cre, only recombining *KP* in AT2 cells^12^) and could confirm that *KP*-Old mice exhibited a reduced tumor burden compared to *KP*-Young mice (Extended Data Fig. 1d).

**Figure 1.**
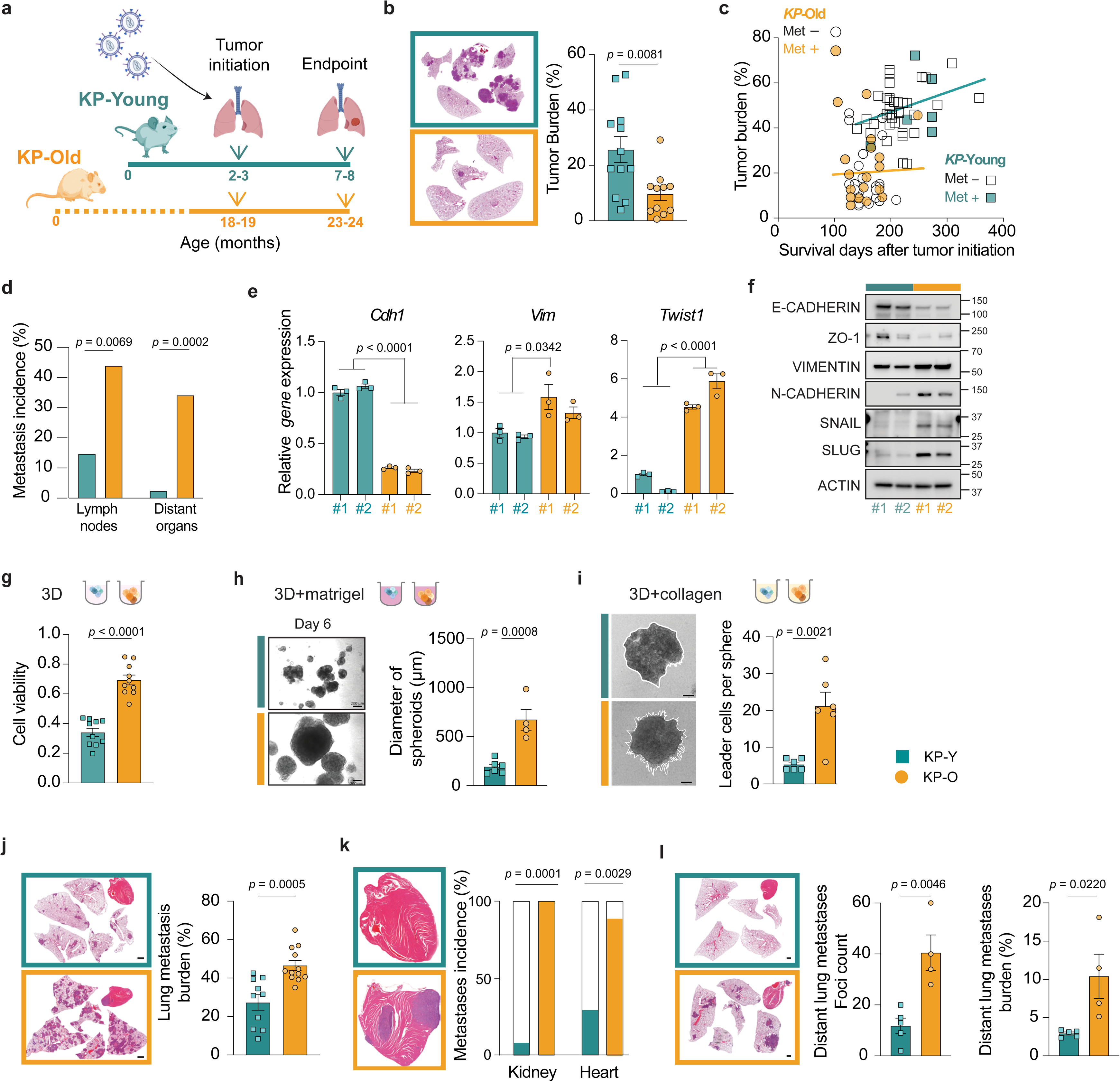
Aging favors lung cancer metastasis over primary tumor growth. **a**.Timeline of the *in vivo* experimental setup. Young and Old *Kras*^LSL-G12D/+^; *Trp53*^flox/flox^ (*KP*) mice were intratracheally instilled with *Cre*-encoding viruses (Lenti-Cre) at 2 and 18 months of age respectively and culled at 7 and 24 months of age respectively. **b.** Left, hematoxylin and eosin (H&E)-stained lung sections of the mice. Right, quantification of lung tumor burden (total tumor area/total lung area) in KP-Young (*n* = 12) and KP-Old (*n* = 11) mice at 21 weeks after tumor initiation induced by Lenti-Cre. **c.** Correlation of primary lung tumor burden with days after Lenti-Cre in KP-Young (*n* = 42, squares, *p* = 0.0731) and KP-Old (*n* = 41, circles, *p* = 0.7026). Mice with metastasis are shown as green squares (KP-Young) and gold circles (KP-Old). **d.** Local lymph nodes and distant metastases incidence of the same mice as in **(c). e**. Relative mRNA expression of epithelial mesenchymal transition (EMT) markers *Cdh1 (*E-CADHERIN), *Vim* (VIMENTIN) and *Twist1* (TWIST) (*n* = 3) in KP-Y and KP-O primary cultures. **f.** Western blot analysis of indicated EMT markers in KP-Y and KP-O cultures with ACTIN as a loading control (*n* = 2). **g**. KP-Y and KP-O primary cultures were seeded in ultra-low attachment plate (ULA) and their relative cell viability was measured 48 h after seeding (*n* = 10). **h**. KP-Y and KP-O primary cultures seeded in ULA plates with 5% Matrigel. Left, representative brightfield microscopic images on day 6 are shown. (Scale bar, 200 μm). Right, quantification of the diameter of spheroids formed by KP-Y (*n* = 6) and KP-O (*n* = 4) measured on day 6. **i.** For invasion assay, KP-Y and KP-O primary cultures were seeded into a collagen matrix in ULA plates. Left, representative brightfield microscopic images 48 h after seeding are shown. Right, quantification of leader cells (independent invasive structures) per spheroid formed by KP-Y (*n* = 6) and KP-O (*n* = 6). (Scale bar, 200 μm). **j-k.** KP-Y (*n* = 10) and KP-O (*n* = 12) primary cultures were injected intravenously in mice. **j**. Left, H&E-stained lung and heart sections of the mice 21 days after injection. Right, quantification of lung metastasis burden (total tumor area/total area). (Scale bar, 1000 µm). **k**. Left, H&E-stained heart sections of the mice (Scale bar, 1000 µm). Right, quantification of metastasis incidence in the kidney and heart of the same mice as in **(j)** 21 days after injection. **l.** KP-Y (*n* = 5) and KP-O (*n* = 4) primary cultures were injected subcutaneously in mice, and distant lung metastases analyzed 28 days after injection. Left, H&E-stained lung and heart sections of the mice. Middle, lung foci number. Right, lung metastasis area. Data presented as mean values ± s.e.m. Statistical significance was assessed by two-tailed unpaired t-test **(b, g, h, i, j,l)**; Simple linear regression **(c)**; Ordinary one-way ANOVA with Tukey’s multiple comparisons **(e)** two-sided Chi-square test **(d, k)**.

We next investigated whether the reduced tumor burden observed in *KP*-Old mice could be seen in additional time points than at 24 months. We found that primary lung tumor burden remained low across all time points measured in *KP*-Old mice (106-247 days) while in *KP*-Young mice we observed increasing tumor burden over time (140-357 days) (Fig. 1c and Extended Data Fig. 1e) and increased survival (Extended Data Fig. 1f, g). Strikingly, in contrast to our expectation given the low primary lung tumor burden, *KP*-Old mice had a marked increase in the incidence of metastasis at local lymph nodes and at distant organs compared to *KP*-Young mice (Fig. 1c, d). To investigate why *KP*-Old mice had more metastasis despite low primary tumor burden, we next established primary tumor cultures (*n* = 2) from *KP*-Young and *KP*-Old mice, hereafter designated *KP*-Y and *KP*-O cultures, respectively (Extended Data Fig. 2a). *KP*-Y and *KP*-O cultures proliferated at the same rate (Extended Data Fig. 2b, c) and expressed similar levels of pro-SPC (Extended Data Fig. 2c), in line with tumors derived from AT2 cells. However, gene and protein expression of epithelialmesenchymal transition (EMT) markers were increased in *KP*-O cultures (Fig. 1e, f), in line with a more metastatic profile. We next seeded the primary tumor cultures in ultra-low attachment plates for assessing spheroid growth in 3D conditions and resistance to anoikis, a form of programmed cell death induced in epithelial cells when detached from the extra-cellular matrix (ECM), both important features during metastatic dissemination. We found that *KP*-O cultures displayed increased cell viability (Fig. 1g and Extended Data Fig. 3a, b), reduced levels of caspase 3/7 activity (Extended Data Fig. 3c, d), and augmented growth over a period of 8 days (Extended Data Fig. 3e) compared to KP-Y cultures. Next, we seeded *KP*-O and *KP*-Y cultures in 3D conditions with Matrigel or Collagen to model ECM-tumor cell interactions. *KP*-O cultures formed larger spheroids in the presence of Matrigel (Fig. 1h and Extended Data Fig. 3f), and they formed more invasive structures in the presence of collagen (Fig. 1i) compared to *KP*-Y cultures. Altogether, this suggests increased metastatic potential in *KP*-O cultures compared to *KP*-Y cultures *in vitro*.

To assess the metastatic potential *in vivo*, we next injected primary *KP*-O and *KP*-Y cultures into the bloodstream of mice through the tail vein to measure their ability to survive in the bloodstream and metastasize to the lungs (Fig. 1j). We found that mice injected with *KP*-O cultures had increased metastasis burden in lungs, and more metastases in kidneys and heart, compared to mice injected with *KP*-Y cultures (Fig. 1j, k). These findings underscore the diverse tropism of *KP*-O cultures but did not inform further about their ability to disseminate and colonize distant sites. Therefore, we next grafted *KP*-O and *KP*-Y cultures subcutaneously in the flank of mice, and quantified distant metastasis. Histological analysis revealed elevated counts of distant lung metastasis foci and increased lung metastasis burden (Fig. 1l) which is in line with increased metastatic capabilities in *KP*-O cultures.

### Aging alters progression of human NSCLC

Next, we assessed the relevance of our findings from *KP-Old* mice to clinical reality of NSCLC patients. We extracted information regarding age, stage at diagnosis, and *KRAS* mutational status from patient charts of all consecutively diagnosed NSCLC patients in Region Västra Götaland and Halland (Western Sweden), between 2016 and 2018 (*n* = 997) (Fig. 2a and Extended Data Table 1). Among them, 368 (36.9%) harbored a mutation in *KRAS* (*KRAS*^MUT^) and 629 patients did not (*KRAS*^WT^). The median age at diagnosis was 71 years for the overall cohort (Extended Data Table 1) and in both mutational groups (Fig. 2b and Extended Data Table 1). We next analyzed this cohort by defining two age groups: less than 60 (< 60) years and old group spanning 65-75 years. The latter group corresponded to the age of *KP*-Old mice at sacrifice (± 5 years), aligned with the median age at diagnosis of the cohort and represented 65.2% of *KRAS*^MUT^ patients diagnosed. We found that the frequency of *KRAS* mutations was higher in the 65-75 aged group compared to the younger group (Fig. 2c), which was also seen in the MSK-IMPACT^13^ dataset (*n* = 435) (Extended Data Fig. 4a). Furthermore, 65-75-year-old *KRAS*^MUT^ patients were more frequently diagnosed with loco-regionally advanced and metastatic disease (Stage IIIb-IV NSCLC) compared to younger < 60-year-old *KRAS*^MUT^ patients, which was not the case for *KRAS*^WT^ patients (Fig. 2d).

**Figure 2.**
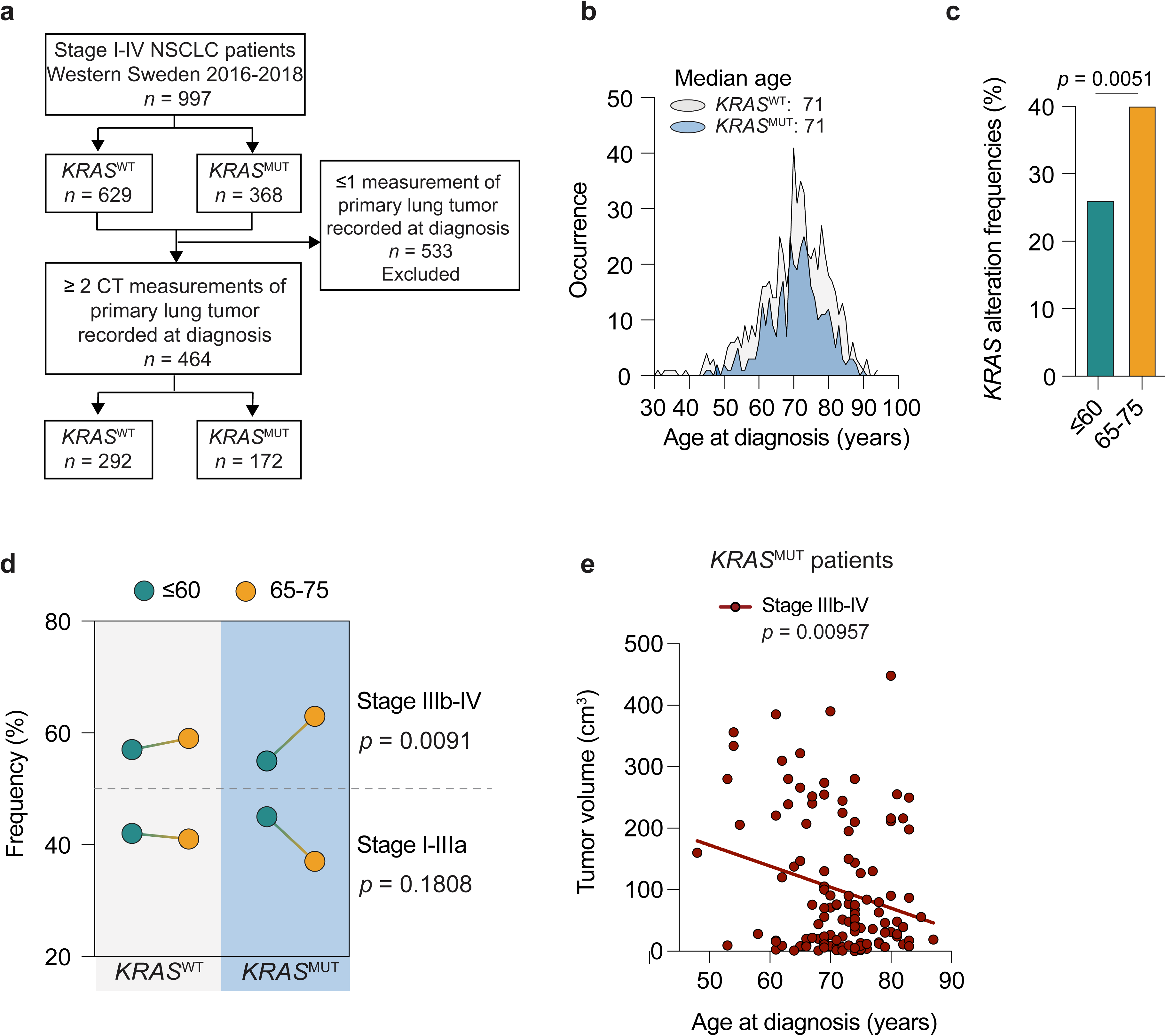
Aging increases metastasis risk despite smaller primary tumor in human NSCLC. **a**. Flow chart showing patient selection for the study. Non-small cell lung cancer (NSCLC); *KRAS* wildtype (*KRAS-* ^WT^), *KRAS* mutated (*KRAS*^MUT^). **b**. Frequency distribution of age at diagnosis for patients harboring a *KRAS*^WT^ (light grey) or *KRAS*^MUT^(blue) alteration. **c.** Left, Incidence of *KRAS* mutations frequencies in young (≤ 60 years) and old (65-75 years) patients at diagnosis in the Western Sweden cohort (≤ 60, *n* = 124; 65-75 years, *n* = 476) **d.** Frequency of young (≤ 60 years, green circles) or old (65-75 years, gold circles) patients from the Western Sweden cohort diagnosed with Stage I-IIIa (*n =* 241) or Stage IIIb-IV (*n =* 358) NSCLC, harboring an alteration in *KRAS (KRAS*^MUT^) or not *(KRAS*^WT^). **e.** Correlation analysis showing negative relation between tumor size and age at diagnosis in Stage IIIb-IV *KRAS*^MUT^ patients (*n* = 117) from the Western Sweden cohort. Statistical significance was assessed by two-sided Fisher’s exact test **(c),** Chi-square test **(d)** and simple linear regression **(e)**.

We next extracted data on primary tumor measurements from patient charts in which ≥ 2 CT-measurements were recorded (*n* = 464) (Fig. 2a). Among them, 172 patients harbored a mutation in *KRAS* (*KRAS*^MUT^) and 292 patients did not (*KRAS*^WT^) (Extended Data Table 2). Importantly, among *KRAS*^MUT^ NSCLC patients with loco-regionally advanced and metastatic disease (Stage IIIb-IV) (*n* = 117) there was a clear decrease in primary tumor size with increasing age at diagnosis (Fig. 2e, *p* = 0.00957), resembling what we observe in metastasis prone *KP*-Old mice (Fig. 1c). We did not observe a similar trend among early stage *KRAS*^MUT^ (Extended Data Fig. 4b) or in *KRAS*^WT^ NSCLC cases across all stages (Extended Data Fig. 4c). Taken together, these findings suggest that increased metastasis incidence despite decreased primary lung tumor burden in *KP*-Old mice reflect the clinical reality for *KRAS*^MUT^ NSCLC patients aged 65-75 years. Along these lines, we found from multiple publicly available online data repositories a positive correlation between doubling time (hours) and age (years) in a panel of human lung cancer cell lines (*n* = 65) (Extended Data Fig. 4d), as well as EMT markers that were increased in *KP*-O cultures (Fig. 1e,f) (Extended Data Fig. 4e, f).

### ATF4 drives aging-induced metastasis

The expression of genes that drive age-related mechanisms in cells is dictated by permissive chromatin states^14–18^. To elucidate the mechanisms driving altered metastatic ability of lung cancer with aging, we performed unbiased gene expression analysis of *KP*-Y and *KP*-O tumor cultures through RNA-seq and observed 2022 differentially expressed genes (DEGs) between the transcriptomes of *KP*-Y and *KP*-O. Next, to investigate the chromatin landscape underlying the differential expression of age-related genes, we performed an unbiased Assay for Transposase-Accessible Chromatin using sequencing (ATAC-seq). Our analysis revealed 27,206 differentially accessible chromatin regions in *KP*-O cultures compared to *KP*-Y. The extensive chromatin remodeling observed suggests that gene expression changes are regulated at the epigenetic level by altering chromatin accessibility landscapes. We next identified in these datasets top enriched transcriptomic and epigenetic pathways identified by gene set enrichment analyses (GSEA) and genomic regions enrichment of annotations tool (GREAT) respectively. Among the top 8 enriched pathways identified through RNA-seq and ATAC-seq, there was overlap between two major themes including EMT and the unfolded protein response (UPR) (Fig. 3a), both highly enriched in *KP*-O cultures (Extended Data Fig. 5a-e, Fig. 3b and Extended Data Fig. 6a). From the UPR-related genes we identified Activating Transcription Factor 4 (ATF4), the master transcriptional effector of the integrated-stress response (ISR)^19^, as one of the top ranked genes (Extended Data Fig. 6a). Accordingly, an ATF4 gene set was differentially expressed compared to *KP*-Y cultures (Fig. 3c). On protein level, ATF4 and its downstream targets, including phosphorylated 4E-binding protein 1 (4EBP1), asparagine synthetase (ASNS), cystine-glutamate antiporter (SLC7A11), C/EBP homologous protein (CHOP) and branched-chain amino acid aminotransferase (BCAT1) were higher in *KP*-O cultures independent of upstream ISR kinases that activate ATF4, including GCN2, PKR, PERK, HRI and EIF2a (Fig. 3d,e)^20^.

**Figure 3.**
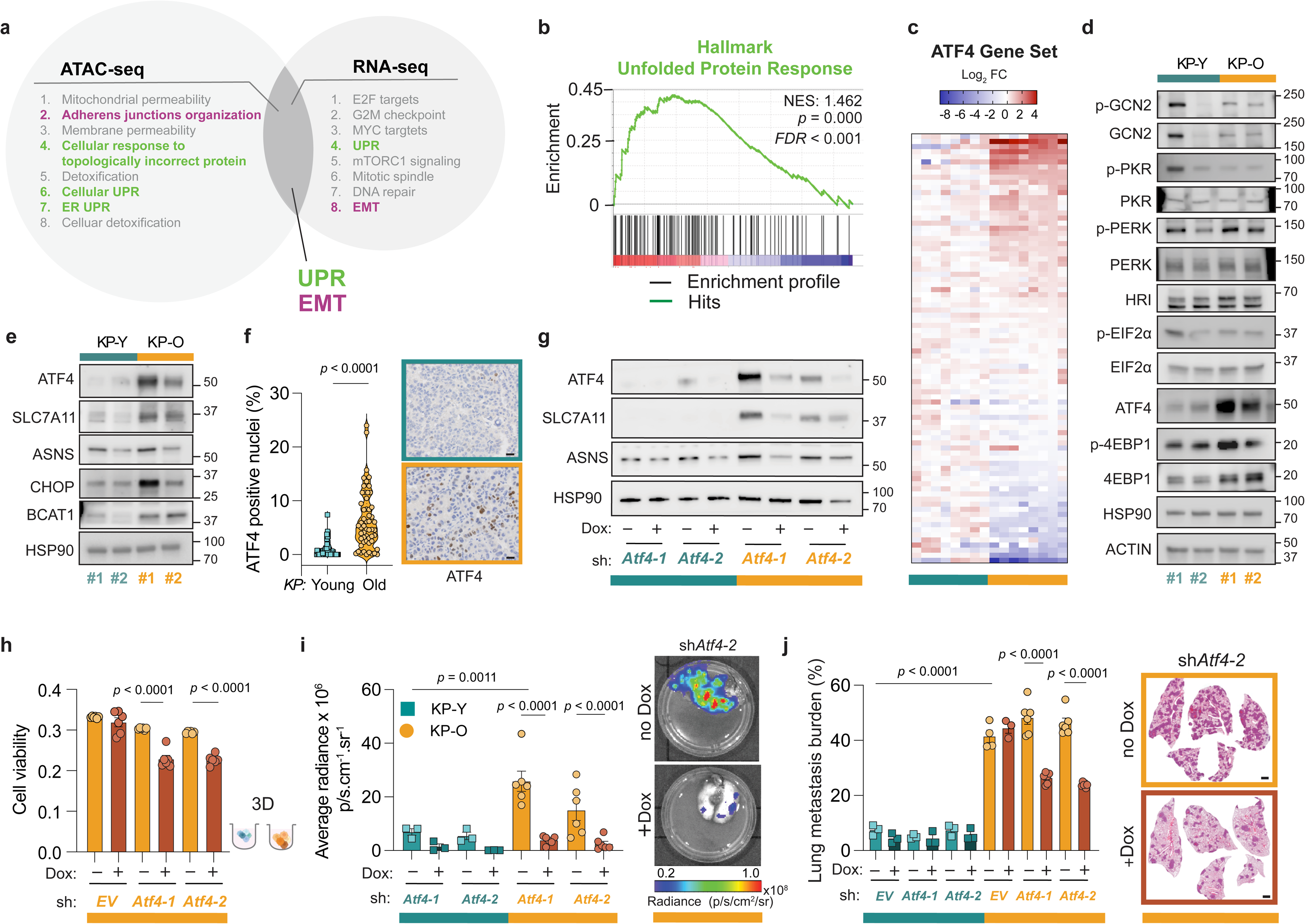
Epigenetic ATF4 induction during aging drives lung cancer metastasis. **a**. Top enriched pathways in KP-O versus KP-Y cultures identified by (left) Genomic Regions Enrichment of Annotations Tool (GREAT) from ATAC-seq and (right) Gene Set Enrichment Analysis (GSEA) from RNA-seq of KP-Y and KP-O cultures. In bold are shown overlapping enriched pathways between both analyses (*n* = 8 per condition). **b.** GSEA enrichment plot showing upregulation of unfolded protein response (UPR) in KP-O compared to KP-Y cultures identified in **(a)**; FDR, false discovery rate; NES, normalized enrichment score. **c.** Heatmap of ATF4 target genes indicating the Log_2_FC expression in KP-Y and KP-O cultures (Human Gene Set: IGARASHI_ATF4_TARGETS_DN; Molecular Signature Database). **d**. Western blot analysis of KP-Y and KP-O cultures showing ATF4 and ATF4 upstream regulators with HSP90 used as loading control. **e**. Western blot analysis of KP-Y and KP-O cultures showing ATF4 and ATF4 downstream targets. HSP90 is used as loading control. **f.** Immunohistochemistry (IHC) staining of ATF4 in lung tumors in KP-Young and KP-Old mice. Left, quantification of ATF4 positive nuclei from lung tumors in KP-Young (*n* = 72 tumors, from 6 mice) and KP-Old (*n* = 55 tumors, from 6 mice) mice after intra-tracheal instillation with Lenti-*Cre-* encoding viruses. Right, representative images of ATF4 IHC staining (Scale bar, 20 µm). **g.** Western blot analysis showing ATF4 and ATF4 targets ASNS and SLC7A11 in KP-Y and KP-O cultures expressing a doxycycline (dox) inducible sh*Atf4*. The sh*Atf4* was induced for 72 h using dox. HSP90 is used as loading control. **h.** KP-O primary cultures expressing a dox inducible sh*Atf4* seeded in ULA plates. Relative viability measured 48 h after seeding. The sh*Atf4* was induced for 72 h using dox before seeding (*n* = 6). (**i-j)**. Intravenous injections of dox-inducible sh*Atf4* in KP-Y (*n* = 3) and KP-O (*n* = 3-6) primary cultures expressing a GFP-luciferase reporter in mice, dox (1 mg/mL) was added to the drinking water. **i**. Left, quantification of lung metastasis burden as average radiance in the lungs 12 days after injection. Right, representative lung ex vivo In Vivo Imaging System (IVIS) images (Scale bar, radiance, p s^−1^ cm^−2^ sr^−1^). **j**. Left, quantification of lung metastasis burden (total tumor area/total area) 12 days after injection. Right, representative hematoxylin & eosin (H&E)-stained lung sections (Scale bar, 1000 µm). Data presented as mean values ± s.e.m. Statistical significance was assessed by Kolmogorov-Smirnov test (**f**) and Ordinary one-way ANOVA with Tukey’s multiple comparisons test (**h-j**).

ATF4 has been implicated in anoikis resistance and metastasis^21,22^. Accordingly, ATF4 expression strongly correlates with liver metastasis penetrance of human cancer cell lines (Extended Data Fig. 6b). In line with this, the percentage of ATF4 positive nuclei was markedly higher in tumors from *KP*-Old mice compared to *KP*-Young (Fig. 3f). Thus, to test if ATF4 regulates aging-induced metastasis, we next targeted ATF4 by CRISPR/Cas9-mediated deletion sgRNAs directed at *Atf4* (Extended Data Fig. 6c) as well as doxycycline-inducible mirE-based shRNA-mediated depletion of ATF4 (Fig. 3g). Protein levels of ATF4 and downstream-targets, including asparagine synthetase (ASNS), cystine-glutamate antiporter (SLC7A11), phosphorylated 4E-binding protein 1 (4EBP1) and others were lowered upon both shRNA and sgRNA based targeting (Fig. 3g and Extended Data Fig. 6c) in both *KP*-Y and *KP*-O cultures. Of note, ASNS and SLC7A11 expression, but not 4EBP1, strongly correlated with ATF4 expression in human metastatic lung adenocarcinoma (LUAD) cell lines, suggesting that the first two could be involved in ATF4-driven metastasis (Extended Data Fig. 6d,e). Indeed, tumors from *KP*-Old mice stained positive for both ASNS and SLC7A11 (Extended Data Fig. 6f, g). Furthermore, knockout or knockdown of *Atf4* reduced anoikis resistance and fitness in *KP*-O but not *KP*-Y cultures when cultured in 3D conditions (Fig. 3h, Extended Data Fig. 7a,b), with little to no effect on cell viability in 2D conditions (Extended Data Fig. 7c.) To confirm whether the increased metastatic ability of tumor cells from *KP*-O mice was ATF4-dependent *in vivo*, we transplanted *KP*-O and *KP*-Y cultures through tail vein injections into mice with ATF4 levels depleted or deleted. Indeed, mice injected with ATF4-deficient *KP*-O cultures had significantly reduced lung metastasis burden compared to non-targeted *KP*-O controls (Fig. 3i, j and Extended Data Fig. 7d,e). In contrast, mice injected with *KP*-Y cultures displayed very little lung metastasis burden compared to *KP*-O injected mice, regardless of ATF4 targeting (Fig. 3i,j and Extended Data Fig. 7d,e). Overall, these results suggest that ATF4 regulates aging-induced metastatic potential in primary *KP*-O cultures.

### ATF4-dependent metabolic rewiring in aging

ATF4 is a major arm of the ISR pathway regulating cellular metabolism and adaptation to metabolic stress conditions^19^. Stable isotope tracing with [U^13^C]-D-Glucose revealed higher lactate production and less tricarboxylic acid (TCA) cycle anaplerosis in *KP*-O cultures (Extended Data Fig. 8a-d). Using isotope-labeled [1,2-^13^C]-D-Glucose, we confirmed increased pyruvate and lactate production through glycolysis – but not through the pentose phosphate pathway (PPP) known to be overactivated in metastatic cancer cells ^23^ - in *KP*-O cultures (Fig. 4a-c). Glucose anaplerosis into the TCA relies on two paths: the pyruvate dehydrogenase (PDH), the main entry point for glucose carbon into the TCA to produce energy, and the pyruvate carboxylase (PC) pathway to maintain TCA intermediates for biosynthesis^24^. Using tracing with [3-^13^C]-glucose, we observed a significantly low carbon incorporation into TCA via pyruvate carboxylase (PC) in *KP*-O cells (Extended Data Fig. 8e-h). This observation led us to hypothesize that *KP*-O cells might exhibit metabolic plasticity and might rely more heavily on alternative metabolic pathways to maintain their TCA cycle intermediates. Thus, we next performed stable isotope tracing with [U^13^C]-L-glutamine which showed enhanced TCA cycle anaplerosis from glutaminolysis in *KP*-O compared to *KP*-Y cultures (Fig. 4d, e and Extended Data Fig 8i). Importantly, increased glutaminolysis is an ATF4-dependent feature, as the contribution of glutamine to TCA intermediates was markedly reduced in *Atf4-*deficient *KP*-O cultures (Fig. 4f, Extended Data Fig. 8j). Finally, we observed a marked increase in extracellular acidification rates (ECAR) in *KP*-O vs *KP*-Y, without any change in oxygen consumption rates (OCR), in line with increased aerobic glycolysis and glutamine anaplerosis (Fig. 4g). Altogether, these results strongly indicate that *KP*-O cultures are metabolically rewired by ATF4 to favor metastasis.

**Figure 4.**
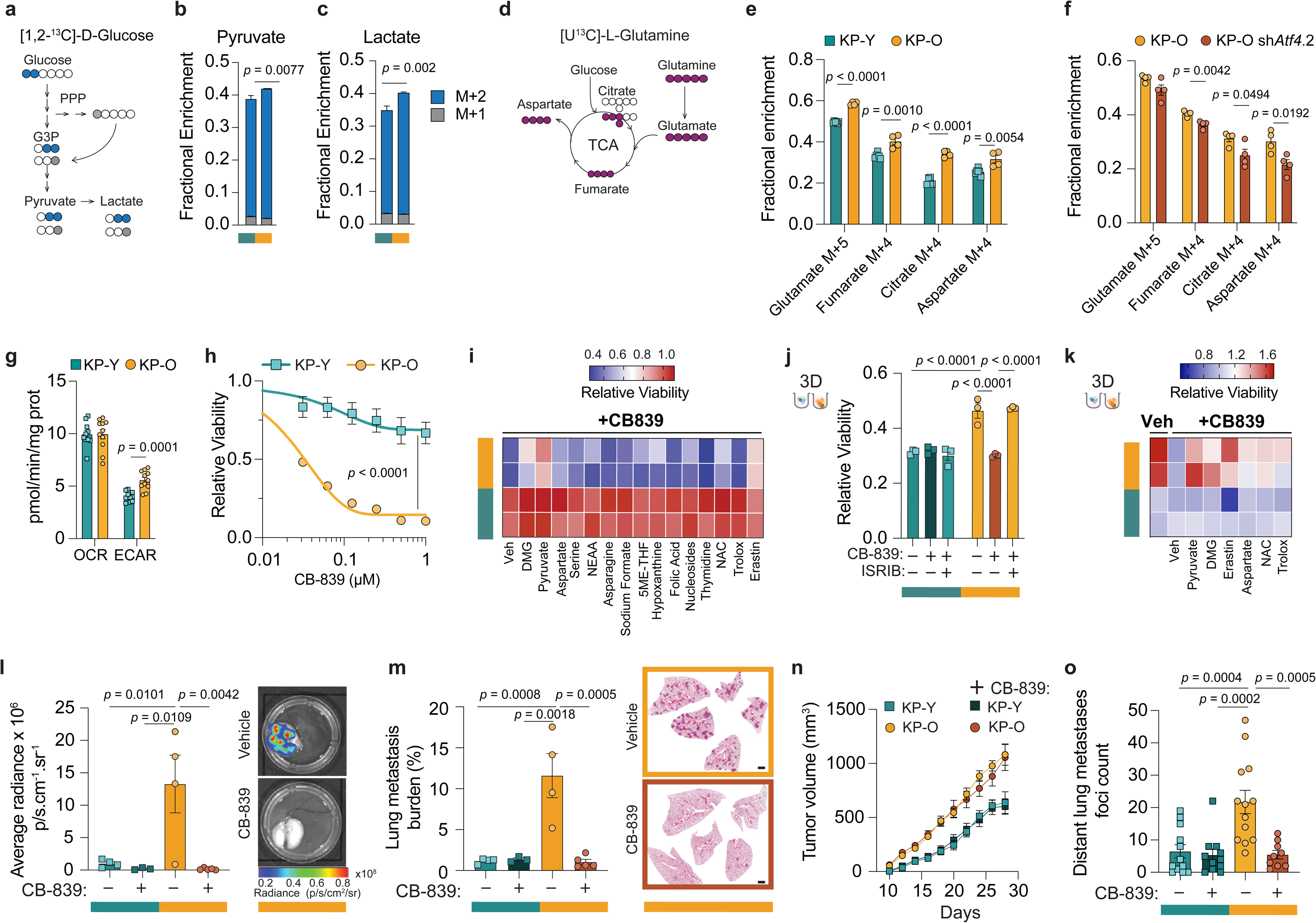
Targeting ATF4-dependent metabolic rewiring blocks aging-induced metastasis. **a.** Schematic depicting the fate of isotope-labeled [1,2-^13^C]-D-Glucose via glycolysis (atoms shown as blue circles) or via the pentose phosphate pathway (PPP) (atoms shown as grey circles). **(b,c).** Fractions of **(b)** pyruvate and **(c)** lactate isotopomers derived from [1,2-^13^C]-D-Glucose in KP-Y and KP-O cultures. (*n* = 6). **d**. Schematic depicting the fate of ^13^C atoms (dark pink circles) from [U^13^C]-L-Glutamine tracing. **e.** Mass isotopomer analysis of TCA cycle intermediates glutamate, fumarate, citrate and aspartate derived from 8 h [U^13^C]-L-Glutamine stable isotope tracing in KP-Y and KP-O cultures (*n* = 6 and *n* =4 respectively). **f.** Mass isotopomer analysis of TCA cycle intermediates glutamate, fumarate, citrate and aspartate derived from 8 h [U^13^C]-L-Glutamine stable isotope tracing in KP-O (*n* = 4) and KP-O sh*Atf4*.2 (*n* = 4). **g**. Seahorse analysis of cellular oxygen consumption rate (OCR) and extracellular acidification rate (ECAR) in KP-O and KP-Y primary cultures (*n* = 10 and 11 respectively). Data are normalized to protein content. **h**. Relative viability assessed by CellTiter-Glo (relative luminescence units) of KP-Y and KP-O cultures after 72 h treatment with CB-839. **i.** Heatmap representing the relative viability values for each indicated primary culture treated with CB-839 and the indicated compounds. All data points are relative to vehicle (veh)-treated controls (*n* = 3). **j**. Relative viability of KP-Y and KP-O cultures in 3D, pretreated for 72 h with 1 µM ISRIB followed by 0.1 µM CB-839 treatment. Cell viability is measured after 48 h of addition of CB-839 (*n* = 3). **k.** Heatmap representing the relative viability values for each indicated primary culture grown in 3D conditions and treated with CB-839 and the indicated compounds. All data points are relative to vehicle-treated controls (*n* = 3). **(l-m)**. Intravenous injections of KP-Y and KP-O primary cultures expressing a GFP-luciferase reporter in mice. Mice were administered 200 mg/kg CB-839 p.o twice/day every other day for the duration of the experiment (n = 3-5 / group). **l.** Left, quantification of lung metastasis burden as average radiance from in the lungs 12 days after injection. Right, representative lung ex vivo In Vivo Imaging System (IVIS) images (Scale bar, radiance, p s^−1^ cm^−2^ sr^−1^). **m.** Left, hematoxylin& eosin (H&E) quantification of lung metastasis burden (total tumor area/total area). Right, representative H&E-stained lung sections (Scale bar, 1000 µm). **(n-o).** Subcutaneous injection of KP-Y and KP-O primary cultures expressing a GFP-luciferase reporter in mice. Mice were administered 200 mg/kg CB-839 p.o twice/day every other day once the tumors reached 100 mm^3^ in size and for the duration of the experiment (*n* = 12, 11, 13, and 9 mice respectively). **n.** Tumor growth of subcutaneous tumors of mice (KP-Y *n* = 24, KP-Y + CB-839 *n* = 22, KP-O *n* = 26 and KP-O + CB-839 *n* = 18 tumors). **o**. H&E quantification of lung metastases foci (*n* = 12, 11, 13, and 9 mice respectively). Data presented as mean values ± s.e.m. Statistical significance was assessed by multiple unpaired t-tests (**b,c,e,f,g),** two-way ANOVA **(h)** and one-way ANOVA with Tukey’s multiple comparisons test **(j,l,m,o)**.

### Targeting metabolic plasticity induced by aging

We next evaluated whether metabolic plasticity in *KP*-O cultures could be exploited for therapeutic targeting (Extended Data Fig. 9a). *KP*-O cultures displayed no sensitivity to pharmacological inhibition of ATF4 by Integrated Stress Response Inhibitor (ISRIB)^25^ (Extended Data Fig. 9b). Furthermore, no differential sensitivity to inhibitors targeting SLC7A11, BCAT or PHGDH was observed (Extended Data Fig. 9c-f). Consistent with increased glutamine utilization, *KP*-O cultures were sensitive to glutamine depletion and glutamine analog DON compared to *KP*-Y cultures, but not to glycolysis inhibition by 2-DG (Extended Data Fig. 9g-i). Glutaminase (GLS) catalyzes the conversion of glutamine into glutamate, the rate-limiting step of glutaminolysis (Extended Data Fig. 9a). Importantly, *KP*-O cultures displayed a marked sensitivity to two independent GLS inhibitors, CB-839 (Telaglenastat) and BPTES (bis-2-(5-phenylacetamido-1,3,4-thiadiazol-2-yl) ethyl sulfide) (Fig. 4h and Extended Data Fig. 9j), in line with previous studies linking high SLC7A11 levels with sensitivity to GLS inhibition ^26,27^. Furthermore, *KP*-O cultures were more sensitive than *KP*-Y cultures to V-9302, an antagonist of the main glutamine transporter Alanine Serine Cysteine Transporter 2 (ASCT2) encoded by *Slc1a5* (Extended Data Fig. 9k).

To pinpoint the source of sensitivity to GLS inhibition in *KP*-O cultures, we pretreated *KP*-O and *KP*-Y cultures with various metabolites, amino acids, antioxidants and small-molecules and found that dimethyl 2-oxoglutarate (DMG), a cell-permeable alpha-ketoglutarate (α-KG) analog, or pyruvate, a glucose-derived TCA cycle carbon source, both rescued CB-839 sensitivity in *KP*-O cultures (Fig. 4i). CB-839 sensitivity was rescued by none of the other tested metabolites or inhibitors except Erastin, strongly indicating *KP*-O cultures are highly sensitive to reduced glutamate levels (Fig. 4i and Extended Data Fig 9a). We next tested if *KP*-O sensitivity to GLS inhibition was ATF4 dependent. Indeed, *KP*-O cultures pretreated with ISRIB, or genetic loss of ATF4 were rescued out of CB-839 sensitivity (Extended Data Fig. 10 a,b). Similarly, *KP*-O spheroid cultures grown in 3D lost their anoikis resistance following glutaminase inhibition with CB-839, another feature that was ATF4 dependent as it was rescued by ISRIB (Fig. 4j). Furthermore, CB-839-induced loss of anoikis resistance in *KP*-O was rescued by DMG, pyruvate and Erastin but not other interventions (Fig. 4k). Finally, we found a correlation between ATF4 expression and CB-839 sensitivity in a panel of human LUAD cell lines derived from metastasis (*n* = 22) but not primary tumors (*n* = 23), suggesting that ATF4 regulates glutamine dependence also in human metastatic LUAD cells (Extended Data Fig. 10c).

To test the *in vivo* relevance of our findings, we next injected *KP*-O and *KP*-Y cultures intravenously or transplanted them subcutaneously into mice and randomized them to CB-839 or vehicle treatment. Following intravenous injection, *KP*-O cultures formed numerous lung metastases in vehicle treated animals but were nearly undetectable in CB-839 treated animals (Fig. 4 l-m). In contrast, *KP*-Y injected animals displayed very little to no metastasis, regardless of treatment (Fig. 4l-m). Following 10 days after subcutaneous transplantation of *KP*-O and *KP*-Y cultures, animals were randomized into Vehicle or CB-839 treatment groups. Interestingly, *KP*-O and *KP*-Y tumors grew at similar speeds regardless of CB-839 treatment (Fig. 4n). However, although *KP*-O tumors were significantly larger than *KP*-Y tumors, there was no difference in final tumor weights between vehicle and CB-839 treated groups (Extended Data Fig. 10d). Strikingly, CB-839 treatment almost completely abrogated distant metastasis of *KP*-O tumors, indicated by distant metastasis foci counts and lung metastasis burden, without impacting the growth or final size of *KP*-O tumors (Fig. 4n-o, Extended Data Fig. 10d-f). In contrast, *KP*-Y tumors and metastasis were unaffected by GLS inhibition (Fig. 4n-o, Extended Data Fig. 10d-f). Our study suggests that aging-induced ATF4-mediated metabolic rewiring constitutes a druggable vulnerability that could be harnessed as a potential adjuvant therapy target for the elderly with NSCLC.

## Discussion

Here, we investigate lung cancer through the lens of aging, uncovering valuable insights into age-induced alterations in tumor progression, tumor metabolism and metastatic potential.

Using an aged *Kras-*driven mouse model for lung cancer and a large and well-characterized clinical cohort of *KRAS* mutant and wild-type NSCLC patients from Western Sweden, our study reveals that the elderly experience reduced primary tumor growth but increased risk of metastasis. Our research reveals that as individuals age, epigenetic changes upregulate EMT, UPR and ATF4, potentially contributing to a more metastatic phenotype. This underscores a critical need for considering age in the personalization of cancer treatments. Based on our findings, there may be indications for initiating aggressive therapy earlier in the form of more frequent neoadjuvant or adjuvant therapy in elderly NSCLC patients to mitigate their increased risk of progressing to metastatic disease.

In parallel and independently of primary tumor expansion, metastatic dissemination can occur early in the course of disease ^28–36^. Reinforcing this non-linear model, our findings reveal that aging stimulates metastasis formation despite a low primary tumor burden. This suggests that while metastasis could be predominantly a late event in younger subjects, it tends to manifest earlier in the elderly. This highlights the importance of early detection and intervention strategies to prevent metastatic disease, which remains the most common cause of cancer-related death^37^. Indeed, managing NSCLC is further complicated by high rates of clinical relapse with disseminated disease, even following curative surgical resection^38^. Therefore, the clinical reality of early dissemination and metastatic dormancy calls for novel therapeutic approaches and earlier interventions among elderly patients diagnosed with NSCLC.

Efficient metastatic cells adapt their nutrient sources, favoring dissemination at the expense of proliferation ^39–45^. Our findings suggest that the relationship between age and lung cancer progression may reflect an integration of age-induced metabolic alterations and the activation of stress pathways that support metastasis. Central to our findings is identifying ATF4, a transcription factor that mediates the ISR and UPR, as a key player in age-induced metastasis. We show that ATF4-dependent glutamine metabolism is essential for metastatic cell survival. Inhibiting glutaminolysis with compounds like CB-839 holds promise as an anti-metastatic or adjuvant therapeutic approach ^46^. However, glutaminase inhibitors such as CB-839 and other novel cancer treatments have often succeeded in pre-clinical trials using young animals but struggled to achieve clinical success. Our findings suggest that age-related metabolic differences in cancer cells may explain this discrepancy. Thus, testing CB-839 as neoadjuvant or adjuvant therapy, while considering age-related metabolic differences between adolescent experimental animals and the average 70-year-old NSCLC patient, could potentially lead to more successful outcomes.

Aging induces significant structural and functional changes in the respiratory system^47^ including alterations in the cell composition of the lung epithelium and the pool of progenitor cells ^48–50^. Specifically, aging leads to epigenetically rooted deregulation and exhaustion of AT2 progenitor cells, the primary cell-of-origin in lung adenocarcinoma^49^. However, in our experiments using KP mice, age-related changes in cell composition or lung progenitors did not influence tumor development: induction of lung tumors with vectors selective to AT2 cells (Adeno-SPC-Cre) or not (Lenti-Cre) both resulted in decreased primary tumor growth, increased metastasis and shorter overall survival in aged animals. Notably, all tumors in both age groups were positive for pro-SPC, indicating consistency in the cell type of origin. The reduced regenerative capacity previously observed in aging AT2 cells ^50^ could nevertheless explain the observed reductions in tumor growth and proliferative capacities of old tumor cells.

This study, together with the lack of comprehensive metastasis data in clinical records, highlights the need for systematic screening protocols to accurately evaluate metastatic burden and guide treatment decisions. Looking ahead, integrating age-related considerations into precision medicine approaches holds promise for improving outcomes in individuals diagnosed with lung cancer.

## Methods

### Mice and *in vivo* studies

*Kras*^LSL-G12D/+^ *Trp53*^flox/flox^ mice (designated as *KP*)^52^ were maintained on a mixed C57BL/6-129/Sv 129 genetic background. Lung tumors were induced in young *KP* (2-3 months old; referred to as *KP*-Young) and old *KP* (18-19 months old, referred to as *KP*-Old) mice through intratracheal instillation with Lenti-Cre as described in ^53^ or with Ad5mSPC-Cre viral particles (PFU 2x10^^7^; University of Iowa; VVC-Berns-1168) under general anesthesia as described in^3^.

*NXG* mice (NOD-Prkdc^scid^ -IL2rg^Tm1^/ Rj) were obtained from Janvier Labs and were 6-10 weeks old at the start of the experiments. For subcutaneous implantations, a total of 2.5 × 10^5^ of GFP-luciferase expressing versions of the indicated cells suspended in PBS were subcutaneously injected into the lower right and lower left flanks of *NXG* mice. Tumors were measured every other day by calipers and the tumor volume was calculated using the formula volume = (length × width^2^) ÷ 2. For intravenous injections, a total of 5 × 10^4^ of the indicated cells were injected into the lateral tail vein of *NXG* mice.

For CB-839 studies, mice were randomized and subjected to treatment with either 200 mg/kg body weight CB-839 (#HY-12248, MedChemExpress) or vehicle. Treatments were administered twice a day every other day following either the tumor-establishment phase (subcutaneous implantations) or within 6 h after the bloodstream injection of cells. The vehicle control consisted of 25% (w/v) 2-hydroxypropyl-β-cyclodextrin in 10 mM citrate buffer (pH 2.0).

For doxycycline-induced *Atf4* silencing, 1 mg/mL of doxycycline (#HY-N0565B, MedChem Express) was administered in the drinking water supplemented with 5% sucrose. Same-sex mice were housed in individually ventilated cages, under a 12 h–12 h light-dark cycle, with temperature and humidity control, enrichment material, and ab libitum rodent chow and water. All mouse experiments described in this study were approved by the Research Animal Ethics Committee in Gothenburg (#2071/19; 2077/19 and 6057/24).

### IVIS imaging

Mice were injected intraperitoneally with 30 mg/mL of D-luciferin (#15225733, FisherScientific) and organs were imaged *ex vivo* 10 min after injection. Luminescence was quantified as radiance with IVIS® Lumina XR series III, for which imaging settings and time were kept constant, or as total flux (ps ^−1^). Analysis was performed with Living Image 4.7.4 software maintaining the region of interest over the tissues as a constant size. The radiance (p/s/cm2/sr) from the region of interest was normalized against the background radiance.

### Histology and immunohistochemical analyses

Lungs were perfused through the trachea with PBS, fixed overnight, transferred to 70% ethanol, embedded in paraffin, cut into 5-µm sections, and stained with hematoxylin/eosin. Tumor burden (percent tumor area per lung area) in H&E-stained sections of all five lung lobes was quantified with BioPix iQ software (v 2.1.4). For immunohistochemical analyses (IHC), deparaffinization was followed by epitope retrieval and blocking of endogenous peroxidase activity using H2O2. Sections were then incubated with antibodies listed in Supplementary Table 1. The total number of positively stained cells in tumors was counted and normalized to the tumor area or to the total number of cells.

### Cells

Primary tumor cultures were isolated from lung tumors of *KP*-Young and *KP*-Old mice approximately after 30 weeks following tumor induction. Primary tumor cultures referred to as KP-Y and KP-O, were isolated with the tumor dissociation kit for mouse (#130-096-730, MACS Miltenyi Biotec) and the gentleMACS Octo Dissociator (#130-093-235, MACS Miltenyi Biotec). Cells were maintained in DMEM medium (FisherScientific) supplemented with 0.1% gentamycin (10 mg/mL, FisherScientific), 2% L-Glutamine (200 mM, FisherScientific) and 10% fetal bovine serum (Thermofisher). All cell lines were routinely tested negative for mycoplasma. For the different assays, cells were seeded in RPMI media (FisherScientific) supplemented with 10% FBS; 1% L-Glutamine, and 0.1% gentamicin.

### Proliferation and viability assays

For population doublings assays, KP-Y1, KP-Y2 and KP-O1, KP-O2 cells were seeded in triplicate in 6-well plates, counted three days later, and re-seeded at the same initial density, for a total of 12-15 days. For cell viability assays, cells were plated in a white, opaque 96-well plate with clear bottom at a density of 2.5 x 10^3^ cells/well in RPMI. 24 h after seeding, drugs were added at the indicated concentrations. 72 h after drug addition, cell viability in the presence of all the compounds was assessed by CellTiter-Glo 2.0 (#G9242, Promega)

### Clonogenic Assay

A total of 5000 KP-Y1, KP-Y2, and KP-O1, KP-O2 cells were seeded in a 10cm plate in RPMI media supplemented with either 2mM, 1mM, 0.5mM or 0.25mM concentrations of L-Glutamine for 6 days. The cell culture medium was replaced after 3 days. Colonies were fixed and stained by incubation in PBS containing 0.05% crystal violet, 1% formaldehyde, and 1% methanol for 20 min. The number of colonies was counted with OpenCFU (Geissmann, 2013).

### 3D cultures and anoikis resistance assay

20 x 10^3^ cells were seeded in an ultra-low attachment plate (#10023683, Corning) and in parallel in a normal attachment 96-well plate in 3-6 replicates per cell line. After 48 h of cell seeding, cell viability was assessed by Cell-Titer-Glo 2.0 (#G9242, Promega). Ultra-low attachment counts were normalized to the attached plate measured 16 h after seeding.

### Spheroid Formation and Collagen Invasion Assays

20 x 10^3^ cells were seeded in an ultra-low attachment plate (#10023683, Corning) with 10% Matrigel (#11523550, FisherScientific). For collagen invasion assays, 10 x 10^3^ cells were seeded in an ultra-low attachment plate (#10023683, Corning). The next day, cells were embedded in collagen (#ECM675, Sigma-Aldrich). Pictures were taken 24-48 h later.

### Caspase activity assay

Briefly, 20 x 10^3^ cells were seeded in an ultra-low attachment plate (#10023683, Corning) and in parallel in a normal attachment 96-well plate in 5 replicates per cell line. After 48 h of cell seeding, caspase activity in cells was measured using the commercially available Caspase-Glo 3/7 Assay (#G8090, Promega). Ultra-low attachment counts were normalized to the attached plate measured 16 h after seeding.

### Cell Trace Proliferation Assay

Cells were labeled using CellTrace Proliferation kit (#C34554, ThermoFischer Scientific) according to the manufacturer’s instructions and seeded in low attachment plates to form spheroids. Single-cell suspensions were prepared from spheroids on days 0, 2, 4, 6, and 8, flow cytometry data were collected on Sony ID7000™ Spectral Cell Analyzer and analyzed using Flow Jo software 10.8.1.

### Lentiviral Production and Transduction

Lentiviruses were produced by transfecting 293FT packaging cells (#R70007, Life Technologies) with lentiviral backbone constructs, packaging plasmid psPAX2 (Addgene plasmid #12260), and envelope plasmid pMD2.G (Addgene plasmid #12259) using the X-tremeGENE™ 9 DNA Transfection Reagent (#6365809001, Sigma-Aldrich). Lentiviral supernatants were collected 2 days after transfection. Target cells were transduced once with lentiviruses supplemented with 8 μg/mL polybrene (#TR-1003-G, Sigma-Aldrich) and selected with puromycin (2 μg/mL, #12122530, FischerScientific). Luciferase-GFP expressing vector pLV[Exp]-EGFP:T2A: Hygro-EF1A>Luc2 was generated by VectorBuilder. The GFP expression was validated using flow cytometry. For CRISPR-mediated gene knockout, the LentiCRISPRv2 (Addgene plasmid #52961) vector was digested with BsmBI and ligated with BsmBI-compatible pre-annealed oligonucleotides (Sanjana et al., 2014; Shalem et al., 2014). Expression of target proteins in CRISPR-knockout experiments was evaluated by western blotting 3–5 days after selection. Sequences are provided in Supplementary Table 1.

### shRNA mediated knockdown

Doxycyline-induced knockdown of *Atf4* was achieved by cloning miR-E shRNAs targeting *Atf4* into the LT3GEPIR vector as previously described in detail (Fellmann et al., 2013). Briefly, LT3GEPIR was digested with XhoI and EcoRI, and purified with a gel extraction kit (#28704, Qiagen). Single-stranded ultramers were amplified with forward primer miRE-XhoI and reverse primer mirE-EcoRI. Amplicons were gel purified, digested with XhoI and EcoRI, cleaned up with a PCR purification kit (#28104, Qiagen) and ligated into the cut LT3GEPIR vector with T4 DNA Ligase at a 3:1 insert: vector molar ratio. Vectors were transduced into cells and selected with 2µg/ml puromycin for two days. Knockdown of ATF4 was verified by western blot analysis following 72 h of treatment with 1 µg/mL doxycycline (#D9891, Sigma-Aldrich). Sequences are provided in Supplementary Table 1.

### Western Blotting

Proteins were isolated by using the 2x Laemmli Buffer (#1610737, BioRad) supplemented with β-mercaptoethanol (#M3148, Sigma Aldrich). Samples were subsequently heated at 95°C for 10 min. Proteins were separated on 4–20% Mini-PROTEAN TGX Stain-Free gel (#4568096, BioRad) and then transferred onto a 0.2 µM nitrocellulose membrane (BioRad), incubated with specific primary antibodies listed in Supplementary Table 1. Protein bands were detected using Clarity Western ECL substrate (#1705061, BioRad) with the Amersham ImageQuant 800 Western blot imaging systems (cytiva). List of antibodies used is provided in Supplementary Table 1.

### Real-Time Quantitative PCR

RNA was isolated with the RNeasy® Plus Mini kit (#74136, Qiagen), and cDNA was synthesized using iScript Adv cDNA Kit for RT-qPCR (#1725038, Bio-Rad). Gene expression was analyzed using the PowerUp™ SYBR™ Green Master Mix (#A25777, Thermo Fisher Scientific) on QuantStudio™ 5 Real-Time PCR system (Thermofisher). List of primers are provided in Supplementary Table 1.

### Reagents

Cells attached or in spheroids were treated with indicated drugs at the following concentrations, unless stated otherwise in the figure: vehicle (DMSO #D8418, Sigma Aldrich), 100 nM CB-839 (#5337170001, Sigma-Aldrich), 2 mM Pyruvate (#11501871; FisherScientific); 50 µM Trolox (#648471, Sigma-Aldrich); 2 mM dimethyl-2-oxoglutarate (DMG, #349631 Sigma-Aldrich), 500 µM N-acetyl-L-cysteine (NAC, # A7250, Sigma-Aldrich), 0.3 µM Erastin (#E7781, Sigma-Aldrich); 20 µM 5-Methyltetrahydrofolic acid disodium salt (5ME-THF, #M0132, Sigma-Aldrich); 5 µM Hypoxanthine (#H9636, Sigma-Aldrich), 1X Nucleosides (#ES-008-D, Sigma-Aldrich), 1X Non-Essential Amino Acids (NEAA, #SH30238.01, Nordic Biolabs), 2mM L-Serine (#S4311, Sigma-Aldrich), 2 mM L-Asparagine (#A4159, Sigma-Aldrich), 2 µM Folic acid (#F8758, Sigma-Aldrich), 50 µM Thymidine (#T1895, Sigma-Aldrich), 1mM Sodium formate (#247596, Sigma-Aldrich), 20 µM L-Aspartic Acid (#A7219, Sigma-Aldrich), 1 µM Inhibitor of Integrated Stress Response (ISRIB, #HY-12495, MedChemExpress), bis-2-(5-phenylacetamido-1,3,4-thiadiazol-2-yl) ethyl sulfide (BPTES, #HY-12683, MedChemExpress), V-9302 (#HY-112683, MedChemExpress), ERG240 (#HY-W193545A, MedChemExpress), branched-chain aminotransferase inhibitor (BCATc, #HY-116044, MedChemExpress), NCT-503 (#HY-101966, MedChemExpress), L-6-Diazo-5-oxonorleucine (DON, #HY-108357, MedChemExpress), 2-Deoxy-Glucose (2-DG, Sigma Aldrich).

### Mitochondrial Respiration

Oxygen consumption rate (OCR) and Extra-Cellular Acidification rate (ECAR) experiments were performed using the XF96pro apparatus from Seahorse Bioscience (Agilent). Cells were seeded to ∼90% confluence for each condition in RPMI. The following day, media was completely replaced with RPMI pH 7.4 (#103576-100, Agilent) containing 10 mM glucose and 2 mM glutamine and incubated for 45 min at 37°C in a CO2-free incubator before measurements. Basal and maximal respiration measurements were obtained by performing a mito-stress test (#103015-100, Agilent) following the manufacturer’s instructions. Data were normalized to protein content using Pierce^TM^ BCA Protein Assay Kit (#23225, ThermoScientific).

### Gas Chromatography/Mass Spectrometry (GC/MS) analysis of polar metabolites and stable isotope tracing

For analysis of tumor primary cultures in attached conditions, 2.5 x 10⁵ cells were seeded in 6 well plates containing 2ml of stable isotope RPMI media supplemented with 10% dialyzed FBS with either 10mM [U13C]-D-glucose, 2 mM [U13C]-L-glutamine (Cambridge Isotope Laboratories), 10mM [1,2 -13C]-D-glucose (Sigma Aldrich) or [3-13C]-D-glucose (Sigma Aldrich) for 8 h. Cells were washed 2X in ice-cold saline and then collected by scraping (for attached cells) or by centrifugation (for spheroids) in 200 µL of 80% (v/v) ice-cold methanol containing 1µg/mL norvaline (Sigma-Aldrich). Samples were then vortexed for 10 min at 4°C and then centrifuged at max speed for 10 min. The supernatant was transferred to fresh tubes and then dried in a speed vac. Dried metabolite extracts were then derivatized with 20mL O-methoxyamine-hydrochloride (MOX) reagent (#89803, Sigma-Aldrich) in pyridine (#270407, Sigma-Aldrich) at a concentration of 20 mg/mL for 60 min at 37°C and 20 mL of N-tert-butyldimethylsilyl-N-Methyltrifluoracetamide with 1% tert-Butyldimethylchlorosilane (TBDMS) (#375934, Sigma-Aldrich) for 60 min at 37°C. After derivatization, samples were analyzed by GC/MS using a DB-35ms column (Agilent Technologies) in an Agilent Intuvo gas chromatograph coupled to an Agilent 5977B mass spectrometer. Helium was used as the carrier gas at a flow rate of 1.2mL/min. 1 µL of sample was injected in split mode (split 1:1) at 270°C. After injection, the GC oven was held at 100°C for one min and then increased to 300°C at 3.5°C /min. The oven was then ramped to 320°C at 20°C /min and held for 5 min at 320°C. The MS system operated under electron impact ionization at 70 eV and the MS source and quadrupole were held at 230°C and 150°C respectively the detector was used in scanning mode, and the scanned ion range was 10-650 m/z. Mass isotopomer distributions were determined by integrating the appropriate ion fragments for each metabolite (Lewis et al., 2014) using MATLAB (Mathworks) and an algorithm adapted from Fernandez and colleagues (Fernandez et al., 1996) that corrects for natural abundance. For all data, total or relative metabolite pool sizes are normalized to cell counts for each condition.

### RNA-Sequencing

Total RNAs were extracted with the RNeasy Plus Mini kit (#74136, Qiagen). RNA quality was assessed using the DNF-471 RNA kit on a fragment analyzer (Agilent). RNA sequencing libraries were prepared according to the Smart-seq2 protocol, developed ^54^ with some minor modifications. Reverse transcription followed by preamplification of purified cDNA was performed as described ^55^. Samples were purified using Agencourt AMPure XP beads (BD Bioscience). Purified samples were further used for library preparation with the Nextera XT DNA library preparation kit and Nextera XT index kit v2 (Illumina), according to the manufacturer’s recommendations. Samples were indexed and amplified by adding 15 µl Nextera PCR master mix and 5 µl of each index, i7 and i5, obtaining a reaction volume of 50 µl. Amplification and indexing were performed in a T100 instrument at 95°C for 10 s, 55°C for 30 s, and as a final step an incubation at 72°C for 5 min.

Using Agencourt AMPure XP (BD Bioscience San Jose, CA, USA) samples were again purified by following the manufacturer’s protocol. Concentrations were measured using Qubit dsDNA high sensitivity Assay Kit, on a Qubit 4 fluorometer (Invitrogen, Thermo Fisher Scientific). The quality control and the fragment size distribution were determined using the DNF-474 High sensitivity NGS kit on a Fragment analyzer (Agilent).

Libraries were pooled and sequencing was performed at the Genomics core facility at the University of Gothenburg on a NextSeq 500 instrument (Illumina) using the NextSeq 500/500 High Output Kit v2.5 (150 cycles) and paired-end sequencing (2x75 cycles).

Indexing of the ENSEMBL GRCm39 reference genome as well as alignment of sequencing reads was performed using STAR version 2.7.9a ^56^. Read count of aligned reads was performed with HTSeq version 0.9.1^57^. Before differential expression analysis, genes with an average read count of less than 3 were excluded. Differential expression analysis was performed with DESeq2 version 1.34.0 ^58^ in R version 4.1.2. First, the four replicate samples within each cell line were merged using the collapseReplicates function. Subsequently, the analysis was run comparing the two groups of cell lines, Young and Old. Significantly regulated genes were defined as having at least two-fold regulation with an adjusted p-value (Benjamini-Hochberg) less than or equal to 0.05. Gene set enrichment analysis was done using the GSEA desktop v4.3.3 ^59,60^. Functional annotation of genes by Gene Ontology (GO) and pathway analysis by KEGG were done using DAVID ^61,62^.

### ATAC sequencing

Library construction for ATAC sequencing of paired-end 150-bp-long reads by the Illumina HiSeq was performed at GENEWIZ Azenta Life Sciences (Germany). Briefly, Sequencing adapters and low-quality bases were trimmed using Trimmomatic 0.38. Cleaned reads were next aligned to reference genome mm10 using Bowtie2. Aligned reads were filtered using samtools 1.9 to keep alignments that have a minimum mapping quality of 30. PCR or optical duplicates were marked using Picard 2.18.26 and removed. Before peak calling, reads mapping to mitochondria (mt) were called and filtered, and reads mapping to unplaced contigs were removed. MACS2 2.1.2 was used for peak calling to identify open chromatin regions. Called peaks were filtered for blacklisted regions to mitigate errors due to mappability. Valid peaks from each group were merged and peaks called in at least 66% of samples are kept for downstream analyses. Reads falling beneath peaks were counted in all samples, and these counts were used for differential peak analyses using the R package Diffbind. All sequencing tracks, bigWig files were viewed using the Integrated Genomic Viewer (IGV 2.16.2). Genomic Regions of Enrichment of Annotations Tool (GREAT) analysis was done to identify the enriched pathways from ATAC sequencing.

### Western Sweden patient cohort

All patients in Western Sweden (8 hospitals) diagnosed with NSCLC from the years 2016 to 2018 and molecularly assessed were included (*n* = 997). Age at diagnosis, histology, and tumor size at diagnosis from CT scans were obtained from patient charts. Patients were assessed with NGS for mutational status on DNA from FFPE blocks or cytological smears using the Ion AmpliSeq™ Colon and Lung Cancer Panel v2 from Thermo Fisher Scientific as a part of the diagnostic workup process at the Department of Clinical Pathology at Sahlgrenska University Hospital, assessing hotspot mutations in *EGFR, BRAF, KRAS,* and *NRAS*.

To obtain the most recent and accurate untreated primary tumor size, measurements were collected from the radiology report of computed tomography (CT) performed before a final diagnosis of NSCLC was established. For patients with two or three tumor dimensions available in their charts, tumor volume was calculated using mathematical formulas ^63^. The formula used for tumors with two dimensions is Volume = (width x width x length/2), where width is the lowest measurement and length is the highest measurement. For tumors with three dimensions, the tumor volume was calculated as follows: Volume = width x length x height. Tumor volume sizes were converted to the same unit, cm^3^. Pearson correlation was used to identify the linear correlation between tumor volume and age for *KRAS* mutated patients and *KRAS* wild-type patients. Approval from the Swedish Ethical Review Authority (Dnr 2019-04771) was obtained prior to the commencement of the study. No informed consent was required due to all data presented in a de-identified form according to the Swedish Ethical Review Authority.

### Online Datasets

Publicly available clinical and cancer genomics data was obtained from cBioPortal for Cancer Genomics, an online tool available at http://www.cbioportal.org/, and from DepMap portal available at https://depmap.org/portal.

Normalized mRNA expression values and age at diagnosis of lung adenocarcinoma patients and cancer cell lines were collected from The Cancer Genome Atlas (TCGA-LUAD) dataset (https://www.cancer.gov/tcga), and Cancer Cell Encyclopedia (CCLE) dataset respectively.

Spearman’s *r* was calculated to analyze the correlation of age with EMT marker expression. Information regarding lung adenocarcinoma organotropism was obtained from MSK-IMPACT data ^13^. Patients were divided into two groups according to age at diagnosis and compared for *KRAS* gene alteration frequencies. Population doublings time and age of cell lines were retrieved from the database https://www.cellosaurus.org/

### Statistical analyses

GraphPad Prism9 v9.0 and v10.0 software were used for statistical analyses. *P-*values were determined by a 2-sided student’s *t*-test for all measurements of tumor burden or when comparing 2 groups, unless stated otherwise in the legends. For the contingency tables, Fisher’s exact test or the chi-squared test was used. One-way ANOVA with Tukey’s *post hoc* test was used for comparisons between multiple groups; for analysis between groups over multiple time measurements, two-way ANOVA was used. Figure legends specify the statistical analysis used. *P-*values of <0.05 were considered significant. All data are represented as mean ± SEM unless otherwise stated.

### Sample sizing and collection

No statistical methods were used to predetermine sample size, but at least three samples were used per experimental group and condition. The number of samples is represented in the graphs as one dot per sample. Samples and experimental mice were randomly assigned to experimental groups. Sample collection was also assigned randomly. All *in vitro* experiments were reproduced at least three times and whenever possible automated quantifications were performed using the appropriate software.

## Acknowledgements

We thank members of the Swedish Lung Cancer Registry and the continuous reporting by Swedish healthcare employees and the staff at the animal facility. This work was supported by the Swedish Research Council (2018-02318 and 2022-00971 to VIS, 2021-03138 to CW), the Swedish Cancer Society (23-3062 to VIS, 22-0612FE to CW), the Gothenburg Society of Medicine (2019; 19/889991 to EAE), Assar Gabrielsson Research Foundation (to AAHP, JD, EAE, CW, and VIS), Department of Oncology, Sahlgrenska University Hospital (to EAE), the Swedish Society for Medical Research (2018; S18-034 to VIS), the Knut and Alice Wallenberg Foundation, and the Wallenberg Centre for Molecular and Translational Medicine (to VIS).

## Author contributions

V.I.S. conceptualized the study. V.I.S., C.W. provided funding, supervised and administered the project. A.A.H.P., S.W.A., C.W., V.I.S. designed and performed experiments and analyses. J.D., K.X.A., A.A.H.P., I.A., A.E.E.Z, K.L.G., D.R., M.S., S.W.A., C.W.

V.I.S. performed experiments. E.A.E., S.I.S, A.H. provided and analyzed the Western Sweden cohort. E.J, H.A and A.S provided transcriptomics platform. S.W.A, M.D, A.A.H.P performed and analyzed metabolomics analyses. A.A.H.P, I.A, E.J. conducted bioinformatics analyses.

R.O.B., A.S.H., A.S. provided resources, protocols and scientific advice. A.A.H.P, S.I.S., C.W, assembled figures. A.A.H.P., S.W.A., C.W., S.I.S and V.I.S. wrote and edited the manuscript. All authors read and approved the final version of the manuscript.

## Data availability

Materials are available upon reasonable request from the corresponding author with an appropriate material transfer agreement. RNAseq and ATACseq data will be made publicly available through the Gene Expression Omnibus and accession numbers provided prior to publication of the manuscript.

Supplementary Information is available for this paper.

## Extended Tables

Extended Data Table 1. Baseline characteristics of patients included in the Western Sweden cohort.

Extended Data Table 2. Characteristics of patients having their primary tumor volume calculated.

## Supplementary Information

Supplementary Table 1. Oligonucleotides sequences and antibodies

---

**Extended Data Fig 1.**
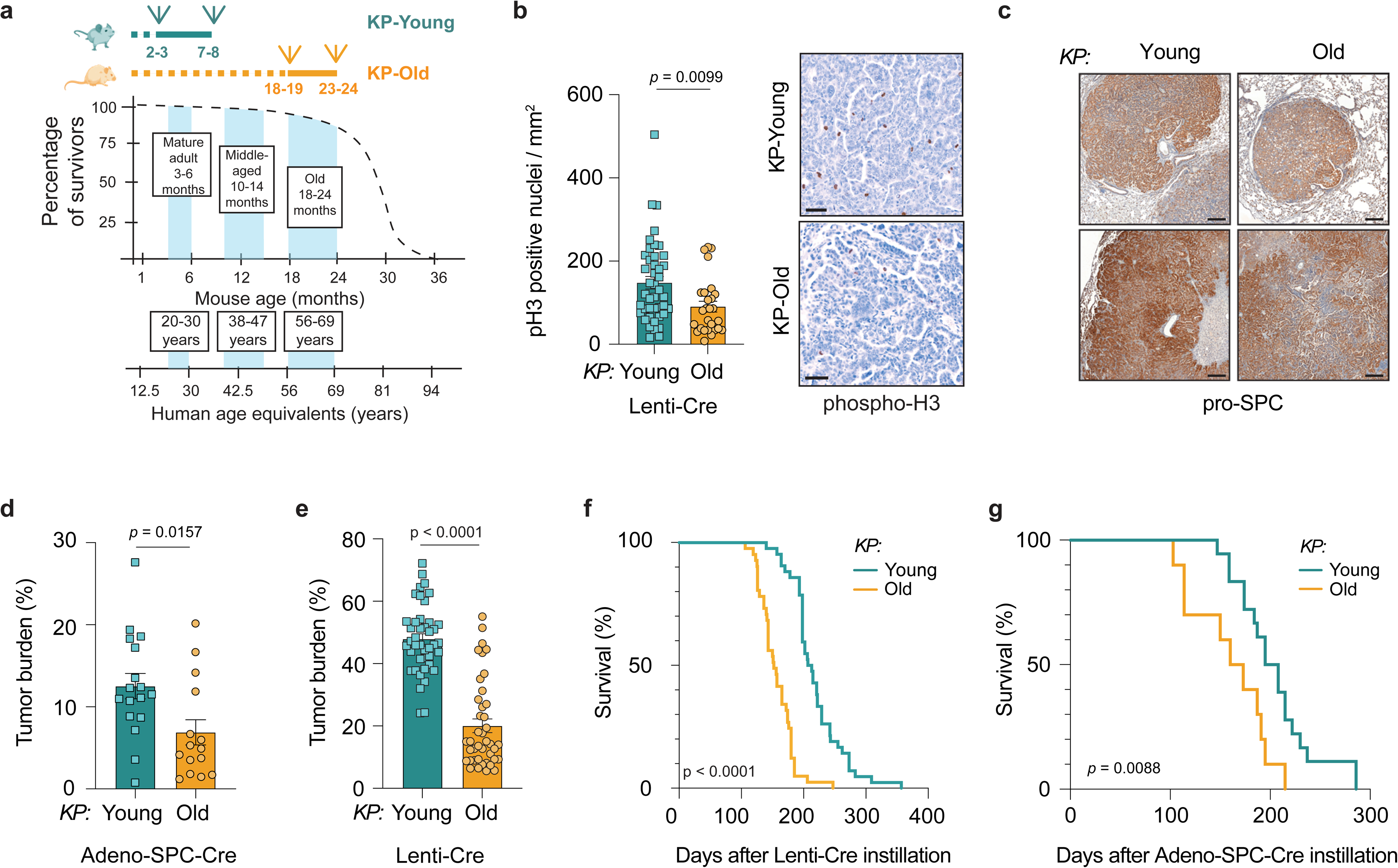
Aging impacts KRAS-driven lung tumor development in mice and their survival. **a.** Schematic depicting the life phase equivalences between mice and humans (adapted from Flurkey et al., 2007). The timeline of the *in vivo* experimental setup is shown on top, indicating tumor initiation and endpoint in KP-Young and KP-Old mice. **b.** Left, quantification of phosphorylated histone H3 (pH3)-positive nuclei per squared millimeter of tumor for assessment of the mitotic index of tumor cells from lung tumors in KP-Young (*n* = 42 tumors, from 6 mice) and KP-Old (*n* = 26 tumors, from 6 mice) mice at 21 weeks after instillation with Lenti-Cre. Right, Representative images from immunohistochemistry staining (IHC) of lung tumors stained for pH3 (Scale bar, 50 µm). **c.** Representative images from IHC stainings of lung tumors stained for alveolar type II (AT2) cell lineage marker Surfactant Protein-C (pro-SPC) in KP-Young and KP-Old 21 weeks after instillation with Lenti-Cre (Scale bar, 200 µm). **d.** Quantification of lung tumor burden (total tumor area/total lung area) in KP-Young (*n* =17) and KP-Old (*n* = 15) mice 17 weeks after instillation with Adeno-SPC-Cre. **e**. Tumor burden of KP-Young (*n* = 42) and KP-Old (*n* = 41) mice after Lenti-Cre instillations from the survival cohort. **f.** Kaplan–Meier survival curves of mice from **(e).** Median survival is 210 and 152 days for KP-Young and KP-Old respectively. **g**. Kaplan–Meier survival curves of KP-Young and KP-Old mice instilled with Adeno-SPC-Cre virus. Median survival is 202 and 167 days for KP-Young (*n* = 18) and KP-Old (*n* = 10) respectively. Data presented as mean values ± s.e.m. Statistical significance was assessed by two-tailed unpaired t-test **(b,d,e)**; Log-rank (Mantel-Cox) test **(f,g).**

**Extended Data Fig 2.**
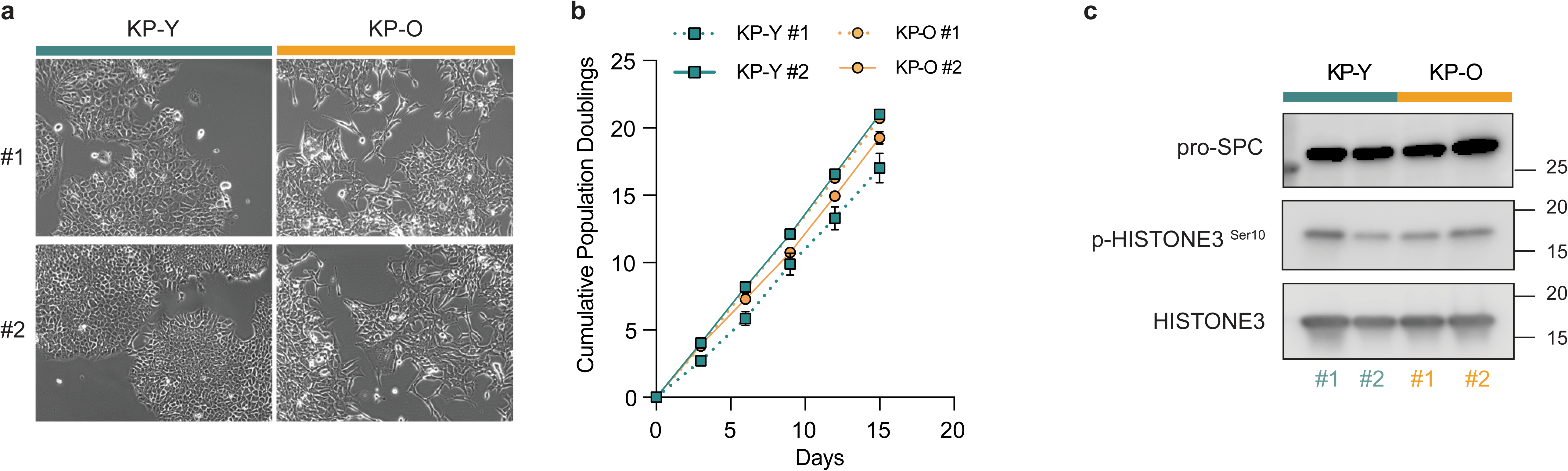
Establishment of primary tumor cultures from KP-Young and KP-Old mice. **a.** Representative brightfield images of primary cultures KP-Y and KP-O established from KP-Young and KP-Old mice respectively (*n* = 2). **b**. Cumulative population doublings of KP-Y and KP-O primary tumor cultures over 15 days (*n* = 2). **c**. Western blot analysis of proliferation marker p-HISTONE3 and pneumocyte type II (AT2 cells) lineage marker Surfactant Protein-C (pro-SPC) in KP-Y and KP-O primary cultures, with HISTONE3 as loading control.

**Extended Data Fig 3.**
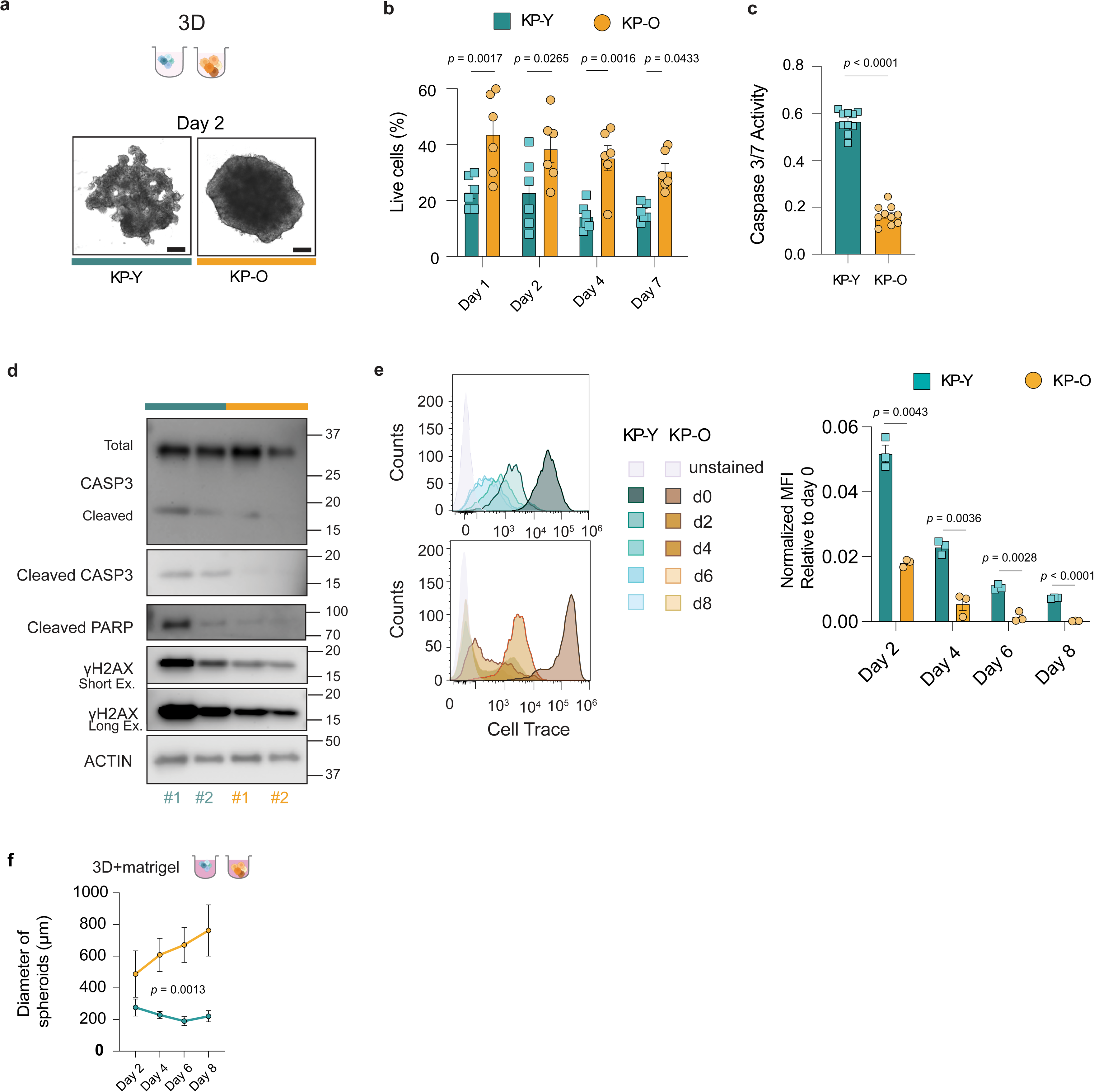
KP-O primary cultures are resistant to anoikis. **a.** Top, Schematic depicting KP-Y and KP-O primary cultures seeded in ultra-low attachment plates (ULA). Bottom, Representative images of KP-Y and KP-O two days after seeding in ULA plates (Scale bar, 200 µm). **b**. KP-Y and KP-O primary cultures were seeded in ULA and viability assessed by trypan blue exclusion. Viability counts shown as percentage of live KP-Y and KP-O cultures after day 1, day 2, day 4 and day 7 (*n* = 6). **c**. KP-Y and KP-O primary cultures were seeded in ULA. Caspase 3/7 activity was measured 48 h later (*n* = 10). **d.** KP-Y and KP-O primary cultures were seeded in ULA for 48 h and western blot analysis was performed with antibodies against CASPASE-3, cleaved CASPASE-3, cleaved PARP, γH2AX short ex: short exposure; long ex: long exposure. ACTIN was used as loading control. **e.** KP-Y and KP-O cultures were labeled with Cell Trace proliferation marker and seeded in ULA plates. Left, representative curves obtained from the CellTrace at day 0, day 2, day 4, day 6, day 8. Right, quantification of Median Fluorescence Intensity (MFI), relative to day 0 (*n* = 3). **f.** KP-Y and KP-O primary cultures seeded in ULA plates with 5% Matrigel. Quantification of the diameter of spheroids formed by KP-Y (*n* = 6) and KP-O (*n* = 4) measured on day 2, day 4, day 6 (shown in Fig. 1h) and day 8. Data presented as mean values ± s.e.m. Statistical significance was assessed by two-way ANOVA **(b**); unpaired t-test **(c)**; multiple unpaired t-test **(e)** and two-way ANOVA (**f**).

**Extended Data Fig 4.**
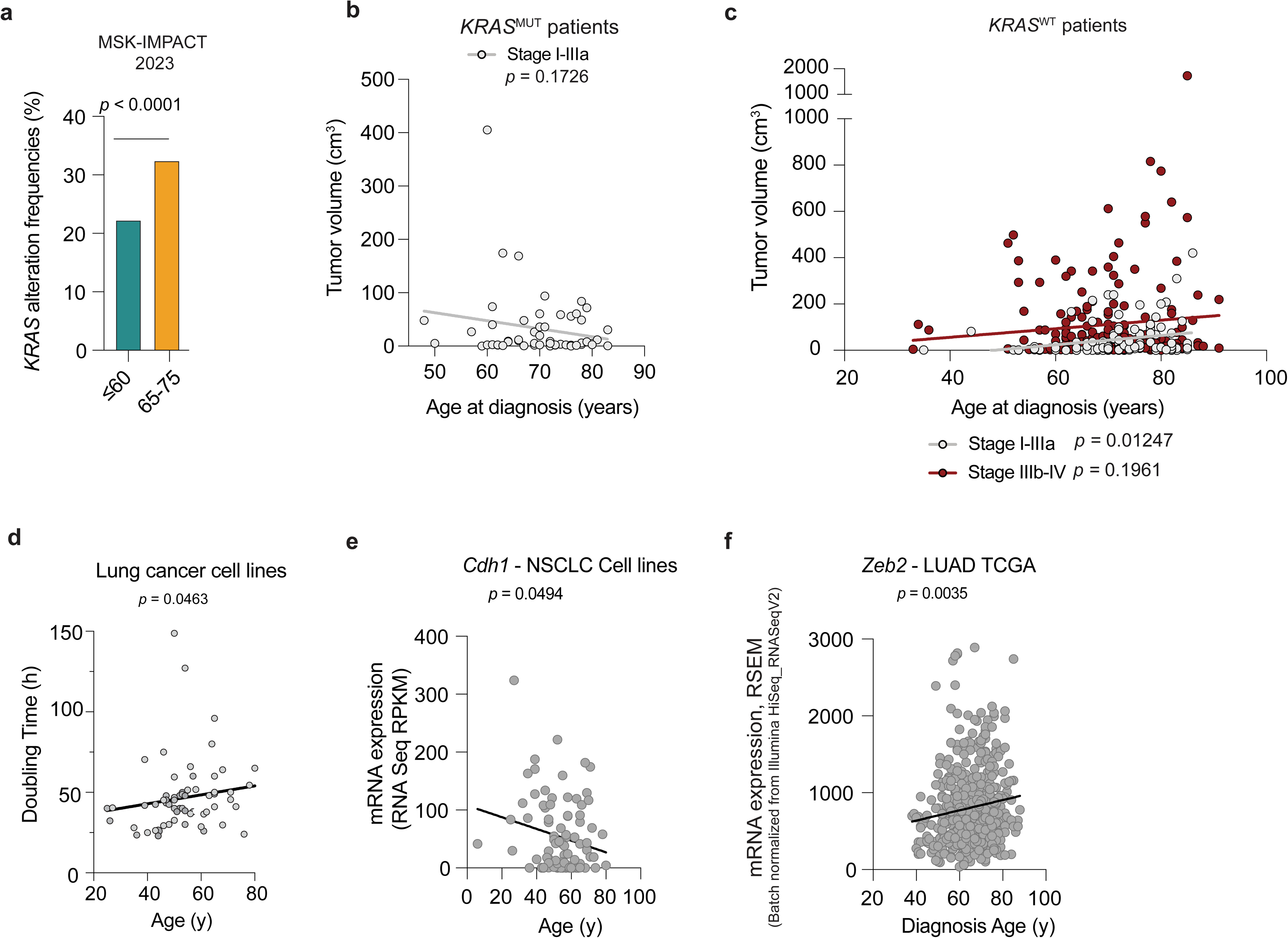
Aging alters tumor volume and metastasis in human NSCLC. **a**. Incidence of *KRAS* mutations frequencies in young (≤ 60 years) and old (65-75 years) patients at diagnosis in the MSK-IMPACT dataset (≤ 60 *n* = 145; 65-75 *n* = 290) (Lengel et al., 2023) **b.** Correlation analysis showing negative relation between tumor size and age at diagnosis in *KRAS*^MUT^ patients Stage I-IIIa from the Western Sweden cohort. (*n* = 55). **c.** Correlation analysis showing relation between tumor size and age at diagnosis in *KRAS*^WT^ patients from the Western Sweden cohort. Stage I-IIIa (*n* = 104); Stage IIIb-IV (*n* = 188). **d.** Analysis showing the positive correlation between doubling time (hours) of lung cancer cell lines and age (years) from EXPASY dataset (*n* = 65). **e**. Correlation analysis of *Cdh1* mRNA expression and age in non-small cell lung cancer (NSCLC) cell lines from c-bioportal database (*n* = 92). **f.** Correlation analysis of *Zeb2* mRNA expression and age at diagnosis in LUAD patients from TCGA database (*n* = 491).Statistical significance was assessed by two-sided Fisher’s exact test **(a),** simple linear regression **(b,c, e, f)** and Spearman correlation **(d).**

**Extended Data Fig 5.**
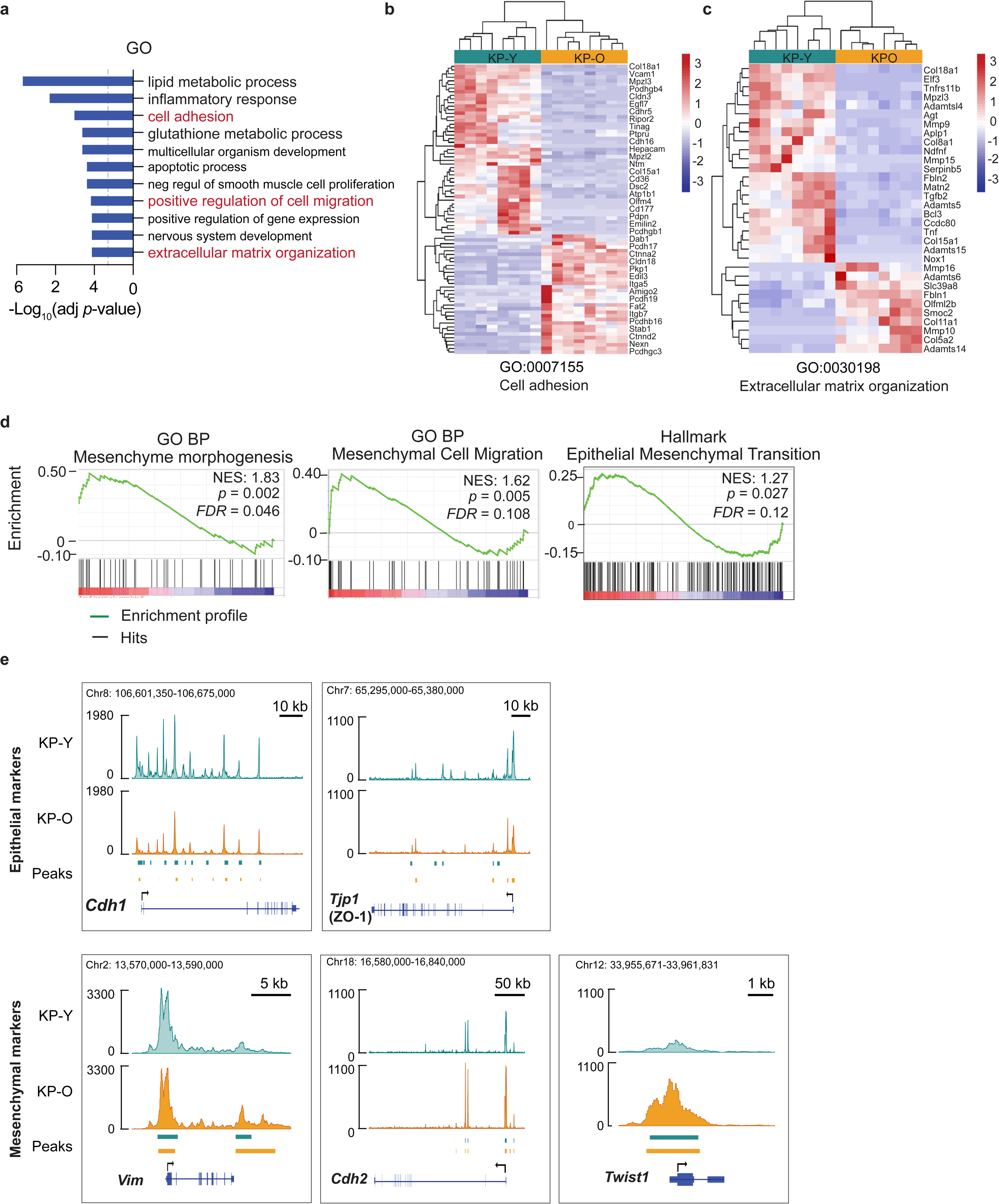
Metastasis-related pathways are enriched in KP-O cultures. **a.** Gene ontology (GO) biological process (BP) terms from enrichment analysis with Database for Annotation, Visualization and Integrated Discovery (DAVID) analysis with the highest statistical significance in KP-O primary cultures compared to that of KP-Y. **b.& c.** Heatmap of normalized counts of genes associated in GO-BP related to **(b)** cell adhesion (every other gene shown) and **(c)** extracellular matrix reorganization in KP-O and KP-Y primary cultures. Expression values for each gene across all samples were normalized by z-score **d.** GSEA enrichment plot showing Mesenchymal morphogenesis, Mesenchymal cell migration and Epithelial mesenchymal transition in KP-O compared to KP-Y cultures; FDR, false discovery rate; NES, normalized enrichment score. **e.** Genome browser view showing the changes in chromatin accessibility of indicated (top) epithelial *Cdh1* (E-CADHERIN) and*Tjp1* (ZO-1) and (bottom) mesenchymal *Vimentin*, *Cdh2* (N-CADHERIN), and *Twist1* markers in KP-Y and KP-O cultures from ATAC-seq. Identified peaks in KP-Y and KP-O are shown below the tracks, in green and orange respectively (*n* = 6).

**Extended Data Fig 6.**
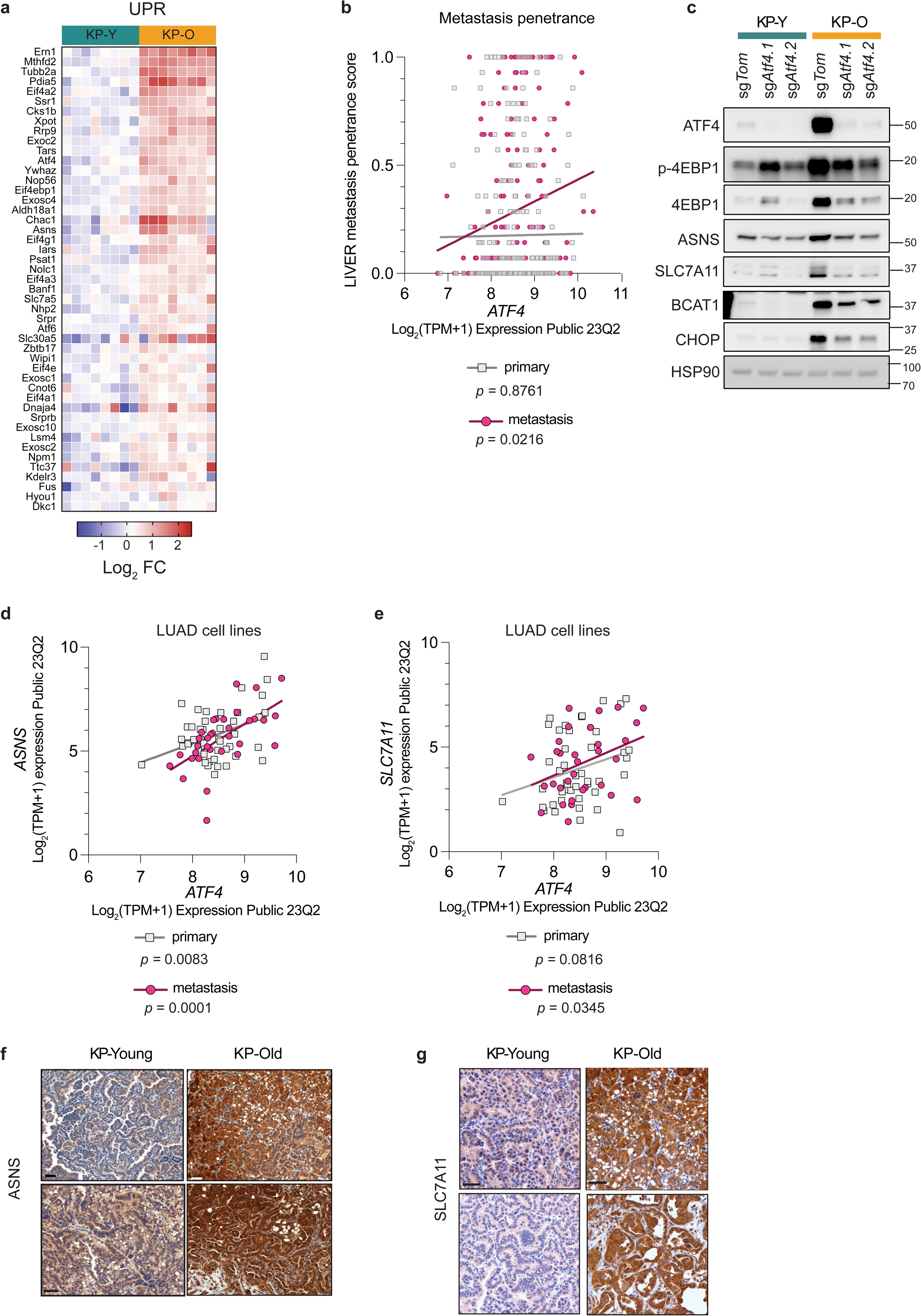
Enrichment of ATF4 pathway with aging in lung tumors and primary cultures. **a.** Heatmap of core enriched genes from UPR pathway identified by GSEA indicating the Log_2_FC expression in KP-Y and KP-O cultures. **b.** Correlation analysis between ATF4 expression and Liver metastasis penetrance in human cell lines from DepMap (*n* = 462). **c.** Western blot analysis showing ATF4 and ATF4 targets in sg*Tom* and Atf4-KO KP-Y and KP-O cultures. HSP90 is used as loading control. **d.** Correlation analysis between *ATF4* expression and ATF4 target gene *ASNS* in human lung adenocarcinoma (LUAD) cell lines Dependency Map (DepMap) portal (*n* = 77). **e.** Correlation analysis between *ATF4* expression and ATF4 target gene *SLC7A11* in human LUAD cell lines from DepMap (*n* = 77). **f.** Immunohistochemistry (IHC) staining of ATF4 target ASNS in lung tumors in KP-Young and KP-Old mice at 21 weeks after intra-tracheal instillation with Lenti-Cre. (Scale bar, 50 µm). **g.** IHC staining of ATF4 target SLC7A11 in lung tumors in KP-Young and KP-Old mice as in (f). (Scale bar, 50 µm). Data presented as mean values ± s.e.m. Statistical significance was assessed by simple linear regression **(b, d, e).**

**Extended Data Fig 7.**
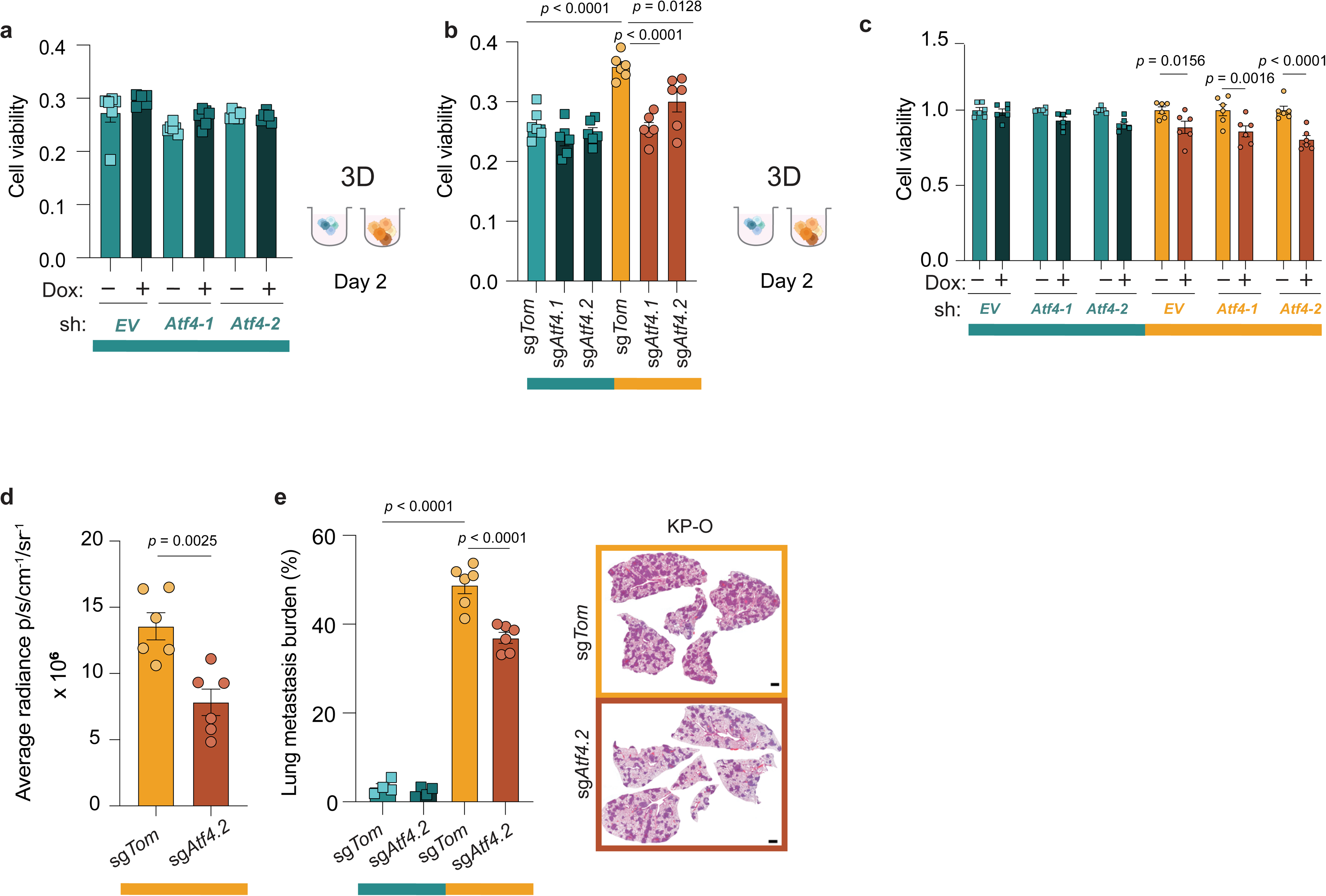
Age-associated metastasis and resistance to anoikis are dependent on ATF4. **a.** KP-Y primary cultures expressing doxycycline (dox) inducible mirE_sh*Atf4* were seeded in ultra-low attachment plates (ULA) plates. Relative viability assessed by Cell-Titer-Glo 48 h later. The sh*Atf4* was induced for 72 h with dox before seeding (*n* = 6). **b.** KP-Y and KP-O expressing either control sg*Tom* or sg*Atf4* were seeded in ULA plates. Relative viability was assessed by CellTiter-Glo (relative luminescence units) 48 h later (*n* = 6). **c**. Relative viability assessed by CellTiter-Glo (relative luminescence units) of KP-Y and KP-O cultures expressing a dox inducible mirE_sh*Atf4* after 72 h of dox induction. All values were normalized to their respective vehicle treated control. (*n* = 6). (**d-e)** KP-Y and KP-O expressing either *sgTom (*KP-Y sg*Tom = 4; K*P-O sg*Tom = 4*) o*r sgAtf4 (KP-Y sgAtf4 = 4; KP-O sg Atf4 = 6) w*ere intravenously injected into mice. **d**. Lung metastasis burden quantified as average radiance in the lungs 14 days after injection**. e.** Left, quantification of lung metastasis burden (total tumor area/total area) 14 days after injection of KP-Y and KP-O primary cultures as described in **(d).** Right, representative hematoxylin and eosin (H&E) -stained lung sections (Scale bar,1000 µm). Data presented as mean values ± s.e.m. Statistical significance was assessed by Ordinary one-way ANOVA with Tukey’s multiple corrections **(b, c, e),** Multiple unpaired t-test **(d).**

**Extended Data Fig 8.**
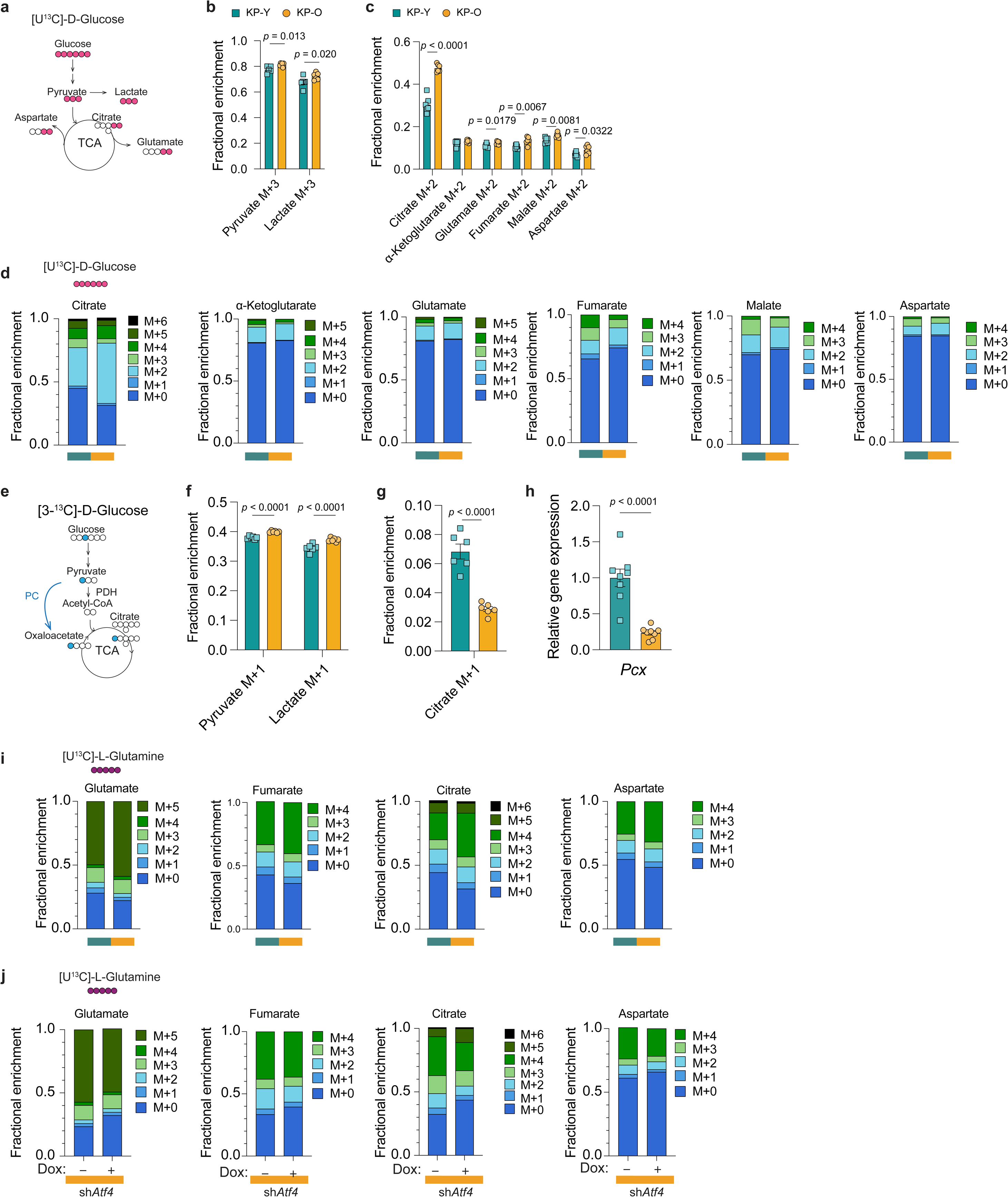
Glucose and glutamine tracing reveals metabolic differences in KP-Y and KP-O cultures. **a**. Schematic depicting the fate of ^13^C atoms (pink circles) from [U^13^C]-D-Glucose tracing. (**b**,**c)**. Mass isotopomer analysis of (**b**) pyruvate and lactate, (**c**) TCA cycles intermediates derived from 8 h [U^13^C]-D-Glucose stable isotope tracing from KP-Y and KP-O cultures (*n* = 6). **d.** Fractions of citrate, α-Ketoglutarate, glutamate, fumarate, malate and aspartate isotopomers derived from [U^13^C]-D-Glucose in KP-Y and KP-O primary cultures (*n* = 6). **e**. A schematic to show the metabolism of isotope-labeled [3-^13^C]-D-Glucose via pyruvate carboxylase (PC) or pyruvate dehydrogenase (PDH) pathways. Blue circles represent ^13^C atoms derived from [3-^13^C]-D-Glucose. **(f-g)**. Mass isotopomer analysis of **(f)** pyruvate and lactate, and **(g)** citrate from 8 h [3-^13^C]-D-Glucose tracing in KP-Y and KP-O (*n* = 6). **h.** *Pcx* gene expression from RNA-seq (*n* = 8). **i.** Fractions of glutamate, fumarate, citrate and aspartate isotopomers derived from [U^13^C]-L-Glutamine in KP-Y and KP-O cultures (*n* = 6 and *n* = 4 respectively). **j.** Fractions of glutamate, fumarate, citrate and aspartate isotopomers derived from [U^13^C]-L-Glutamine in KP-O cultures (*n* = 4). Data presented as mean values ± s.e.m. Statistical significance was assessed by unpaired t-test **(b,c, f-h).**

**Extended Data Fig 9.**
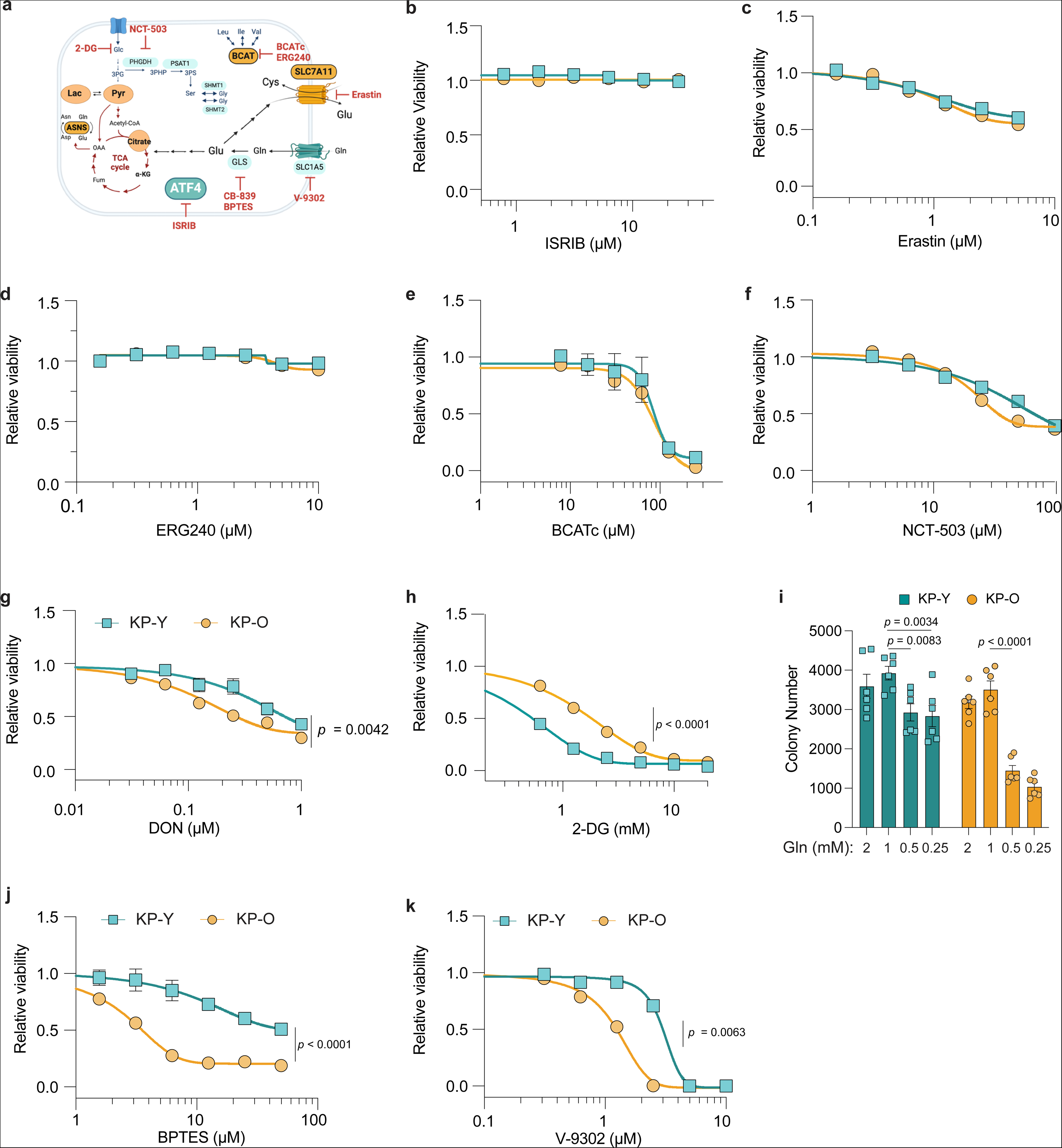
ATF4-directed drug screening identifies sensitivity to glutaminase inhibition in KP-O primary cultures. **a**. Schematics showing the mode of action of the inhibitors used in KP-Y and KP-O cultures. The enzymes and metabolites depicted in orange are ATF4 targets increased in KP-O cultures. **(b-h).** Relative viability assessed by CellTiter-Glo (relative luminescence units) of KP-Y and KP-O cultures after 72 h treatment with **(b)** ISRIB, **(c)** erastin, **(d)** ERG240, **(e)** BCATc, **(f)** NCT-503 and **(g)** DON, **(h)** 2-DG. All values were normalized to their respective vehicle treated control. X-axis depicts respective drug concentration. (*n* = 3). **k.** Colony number of KP-Y and KP-O cultures grown in RPMI media with indicated L-Glutamine (Gln) concentrations for 6 days (*n* = 6). **(j,k)**. Relative viability assessed by CellTiter-Glo (relative luminescence units) of KP-Y and KP-O cultures after 72 h treatment with **(j)** BPTES and **(k)** V-9302. All values were normalized to their respective vehicle treated control. X-axis depicts respective drug concentration. (*n* = 3). Data presented as mean values ± s.e.m. Statistical significance was assessed by by 2-way ANOVA **(b-h,j,k)**, one-way ANOVA with Tukey’s multiple comparisons test **(i).**

**Extended Data Fig 10.**
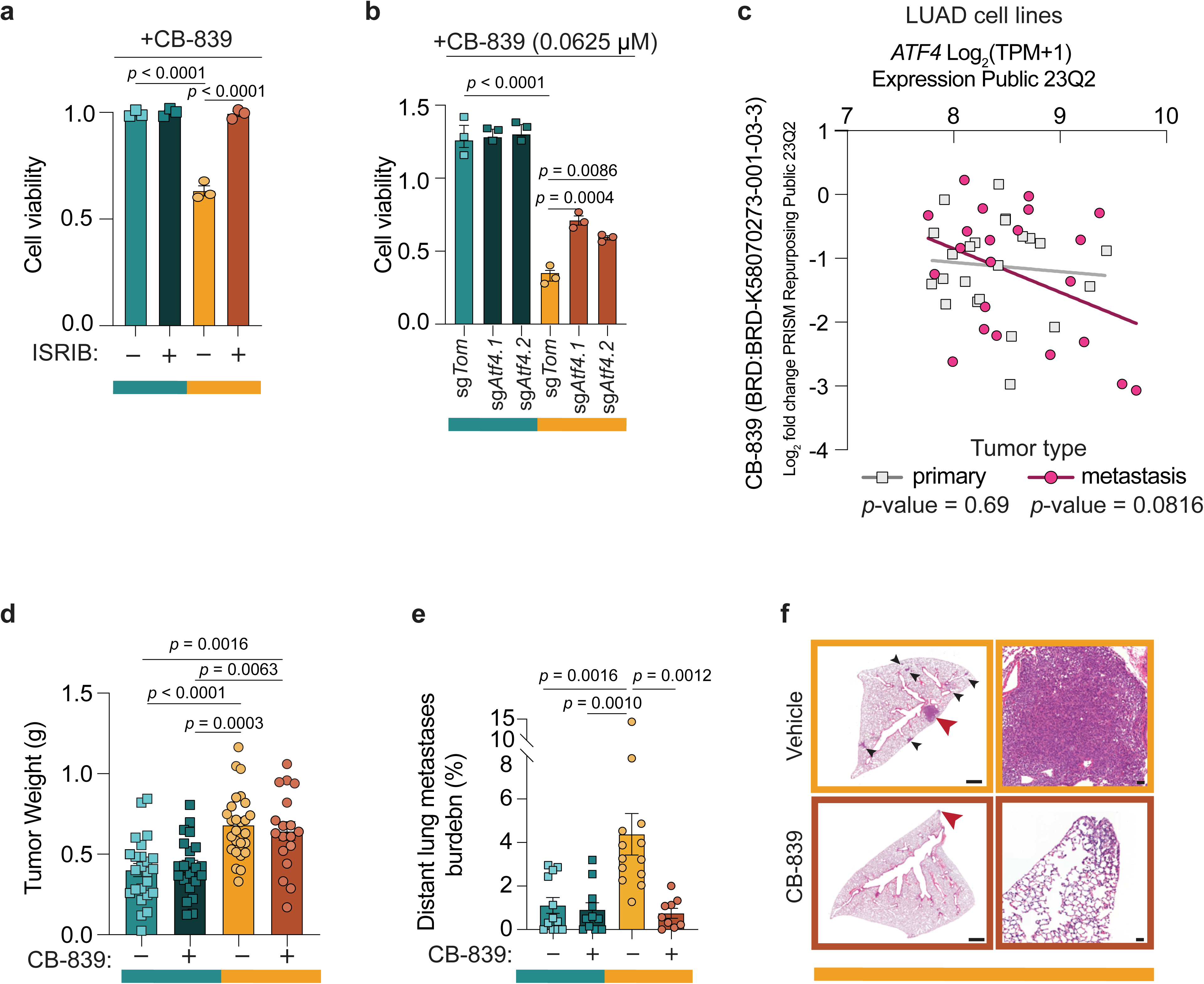
CB-839 sensitivity is dependent on ATF4 in metastatic context. **a.** KP-Y and KP-O primary cultures were pre-treated for 72 h with 1 µM ISRIB followed by 0.1 µM CB-839 treatment for another 72 h. Cell viability was assessed by CellTiter-Glo (relative luminescence units). All values were normalized to their respective vehicle treated control (*n* = 3). **b.** KP-Y and KP-O (sg*Tom* or sg*Atf4*) primary cultures were treated with 0.0625 µM CB-839 for 72 h. Cell viability was assessed by CellTiter-Glo (relative luminescence units). All values were normalized to their respective vehicle treated control (*n* = 3). **c.** Correlation analysis between *ATF4* expression and CB-839 sensitivity in human lung adenocarcinoma (LUAD) cell lines (*n* = 45) from Dependency Map (DepMap) portal. Primary cell lines are depicted with grey squares, metastatic cell lines with pink circles. **(d-f)** KP-Y and KP-O primary cultures were subcutaneously injected in mice. Mice were administered 200 mg/kg CB-839 p.o twice/day every other day for the duration of the experiment once the tumors reached 100 mm^3^ in size. Complementary results presented in Fig. 4 n,o. **d.** Tumor weight of the tumors formed by KP-Y and KP-O primary cultures after subcutaneous injections **e.** Quantification of lung metastasis burden (total tumor area/total area). **f**. Left, hematoxylin and eosin (H&E)-stained lung sections showing metastatic foci in the lungs (black and red arrows). Right, close-up of the area identified by the red arrow in the left panel (Scale bar, 1000 µm and 50 µm). Data presented as mean values ± s.e.m. Statistical significance was assessed by simple linear regression **(c)**, ordinary one-way ANOVA with Tukey’s multiple comparisons test **(a,b,d,e**).

